# Simultaneous GCaMP imaging and focal recording of tonic and phasic synapses: Probing short-term plasticity within a defined microenvironment

**DOI:** 10.1101/2024.08.25.609565

**Authors:** Xiaomin Xing, Chun-Fang Wu

## Abstract

**Key point summary:**

- We developed a protocol for stable simultaneous focal recording and GCaMP imaging on identified motor synaptic boutons in the *Drosophila* neuromuscular preparation with minimized muscular movements in extended concentration ranges of extracellular Ca^2+^ and Sr^2+^.
- This approach directly demonstrated temporal correlation between the dynamics of cytosolic residual Ca^2+^ and activity-dependent synaptic plasticity in tonic and phasic synapses.
- Our data demonstrated presynaptic GCaMP signals markedly lagged behind and poorly correlated with the concurrent transmitter release, but reliably indicated the immediate states of short-term plasticity. GCaMP’s <u>physical-chemical</u> properties allow information extraction on cytosolic Ca^2+^ levels during facilitation and depression phases.
- The decay phase of GCaMP signal often coincided with lingering vesicular releases after stimulation, more pronounced in phasic than tonic synapses and exaggerated in Sr^2+^-containing saline. Lingering releases were coupled with the tendency of asynchronous transmission.

GCaMP fluorescence has been widely used to monitor intracellular Ca^2+^. However, the physiological significance of the GCaMP signal in presynaptic terminals remains to be further elucidated. We investigated how the dynamics of GCaMP signals correlates with the activity dependence of short-term plasticity in synaptic transmission. We devised a local manipulation protocol that minimizes interference from muscular contraction during simultaneous Ca^2+^ imaging and focal recording at the *Drosophila* larval neuromuscular junction (NMJ), where the tonic and phasic excitatory synapses can be compared side-by-side. By confining the local ionic microenvironment, this protocol enabled stable measurements across extended concentration ranges of Ca^2+^ or Sr^2+^ in saline. Compared to tonic synapses, phasic synapses displayed stronger GCaMP signals, along with faster facilitation and more severe depression. Upon repetitive stimulation (40 Hz), facilitation of transmission occurred during or immediately prior to the early rising phase (0.25 s) of the GCaMP signal, which could subsequently convert into a depression phase of transmission decline, most evident during a steeper and longer rise of GCaMP signals in higher Ca^2+^ saline. Typically, deepest depression occurred when GCaMP signals rose to a plateau. Phasic synapses with stronger GCaMP signal and deeper depression, more often exhibited lingering post-stimulation releases. In both tonic and phasic synapses, replacing Ca^2+^ with Sr^2+^ induced extreme asynchronous transmission coupled with post-stimulation lingering releases during the decay of GCaMP signals. Further applications of this focal recording-local manipulation protocol may help to probe additional mechanisms underlying synaptic transmission and plasticity.

## Introduction

Synaptic transmission is a fundamental process that underlies the functioning of the peripheral and central nervous systems. Synapses of different kinetic and activity-dependent properties give rise to the rich variety of neural function and behavioral patterns. The diverse synaptic types across invertebrate and vertebrate species nevertheless share the common scheme of basic cellular and molecular mechanisms. To trigger synaptic transmission, action potentials propagate down the axon to depolarize the presynaptic terminal, which opens the local voltage-gated Ca^2+^ channel (CaV), and subsequent Ca^2+^ influx initiates vesicular exocytosis and transmitter release, which in turn opens the ligand-gated ion channels in the postsynaptic membrane to generate synaptic potentials (Zucker, 1996; Aidley, 1998). Subsequently, the temporal and spatial dynamics of presynaptic residual intracellular Ca^2+^ orchestrates the various signaling pathways that shape activity-dependent transmission plasticity (Yamada and Zucker, 1992; Regehr, 2012; Weingarten et al., 2022).

GCaMP has been a widely used optogenetic indicator for inferring neuronal activities from fluorescent signals reporting binding to cytosolic free-Ca^2+^ (Nakai et al., 2001; Chen et al., 2013). However, the properties of presynaptic GCaMP signal are less extensively characterized (Reiff et al., 2005; Chouhan et al., 2010; Singh et al., 2018). Simultaneous correlation between presynaptic GCaMP signal and neurotransmitter release has shown a significant disparity in time scale, with delays from a few milliseconds in extracellular field excitatory junction potentials (efEJPs) to several tenths of second in local presynaptic GCaMP signals (Xing and Wu, 2018a and b). Thus, the proper interpretation of the GCaMP signal in relation to the elements of presynaptic physiological processes, in particular, how the dynamic cytosolic free-Ca^2+^ levels regulate synaptic plasticity, remain to be further examined.

Synapses displaying different kinetic properties may converge to innervate the same target cell, resulting in a richer variety of transmission patterns. Well-studied examples include some crayfish and *Drosophila* neuromuscular junctions (NMJs), where both tonic (type Ib) and phasic (Is) excitatory synapses coexist (Atwood et al., 1993; Jia et al., 1993; Nguyen et al., 1997; Lnenicka and Keshishian, 2000). This anatomical layout facilitates direct side-by-side comparisons between two types of synapses with distinct characteristics of transmission. In particular, the translucent *Drosophila* larval NMJ preparation is especially suitable for studies combining electrophysiological recording and optical imaging.

*Drosophila* larval NMJs has been used to study the neurogenetic mechanisms for synaptic development, function and plasticity since the 1970s (Jan and Jan, 1976; Anderson et al., 1988; Johansen et a., 1989; Budnik et al., 1990). A substantial body of work has accumulated from different technical advances by using the larval NMJ preparation, including intracellular recording of excitatory junction potentials (EJPs) with the associated motor axon action potentials (Jan et al., 1977; Wu et al., 1978), voltage-clamp measurements of excitatory junction currents (EJCs) and postsynaptic muscle membrane currents (Wu et al., 1983; Warbington et al., 1996; Wang et al., 2000; Yoshihara and Littleton, 2002), focal loose-patch clamping (Kurdyak et al., 1994; Bradacs et al., 1997), and Ca^2+^ imaging with either synthetic or optogenetic Ca^2+^ indicators (Macleod et al., 2002; Millar et al., 2005; Reiff et al., 2005; Lnenicka et al., 2006; Melom et al., 2013).

Here we report results based on a protocol for local microenvironment control of synaptic boutons to enable direct correlation of the presynaptic GCaMP signal with electrophysiological recording of synaptic transmission in extended divalent cation (Ca^2+^and Sr^2+^) concentration ranges. This approach does not rely on the use of glutamate for desensitizing the postsynaptic glutamate receptor channels (Macleod et al., 2002; Lnenicka et al., 2006), or anesthetization but could similarly minimize muscle contraction at physiological or higher Ca^2+^ concentrations to achieve stable efEJP recordings and associated GCaMP Ca^2+^ imaging, yielding comparisons of phasic and tonic synapses functioning at low and high Ca^2+^/Sr^2+^ concentrations. The results demonstrate distinct Ca^2+^-dependent short-term plasticity parameters of tonic and phasic synapses in relation to their GCaMP signal dynamics, and the progression of facilitation-depression transition during a train of transmission activities. Based on the properties of GCaMPs, the dynamics and concentration ranges of cytosolic Ca^2+^ during the different phases of activity-dependent plasticity could be estimated. Replacement of Ca^2+^ with Sr^2+^ further uncovered factors that affect transmitter release synchronicity, indicating that this approach could be readily extended for exploration of additional aspects of synaptic transmission and plasticity using the *Drosophila* larval NMJ. Over all, our results point up the need to secure comprehensive and kinetically resolved baseline information to enable proper interpretation of GCaMP signals in extracting relevant physiological parameters.

## Materials and Methods

### Fly Stocks

All stocks used were maintained at room temperature (22– 24°C) on standard medium (Frankel and Brosseau, 1968). The genotypes used in this study include: UAS-GCaMP lines and neuron-specific Gal4 lines on the same chromosomes were recombined together to form stable stocks. *UAS-GCaMP1.3* was linked with *c164-Gal4* to form +; *c164-Gal4*::*UAS-GCaMP1.3* (hereafter referred to as *c164-GCaMP1*, used in Figures 10). *UAS-GCaMP6f* was also recombined with *c164-Gal4* to form +; *c164-Gal4*::*UAS-GCaMP6f* (*c164-GCaMP6f*, used in Figures 1, 2). Likewise, +; +; *nSyb-Gal4*::*UAS-GCaMP6m* (*nSyb-GCaMP6m*, used in all the other figures) was constructed. The sources of all the optogenetic lines have been described previously (Xing and Wu, 2018a).

**Figure 1.**
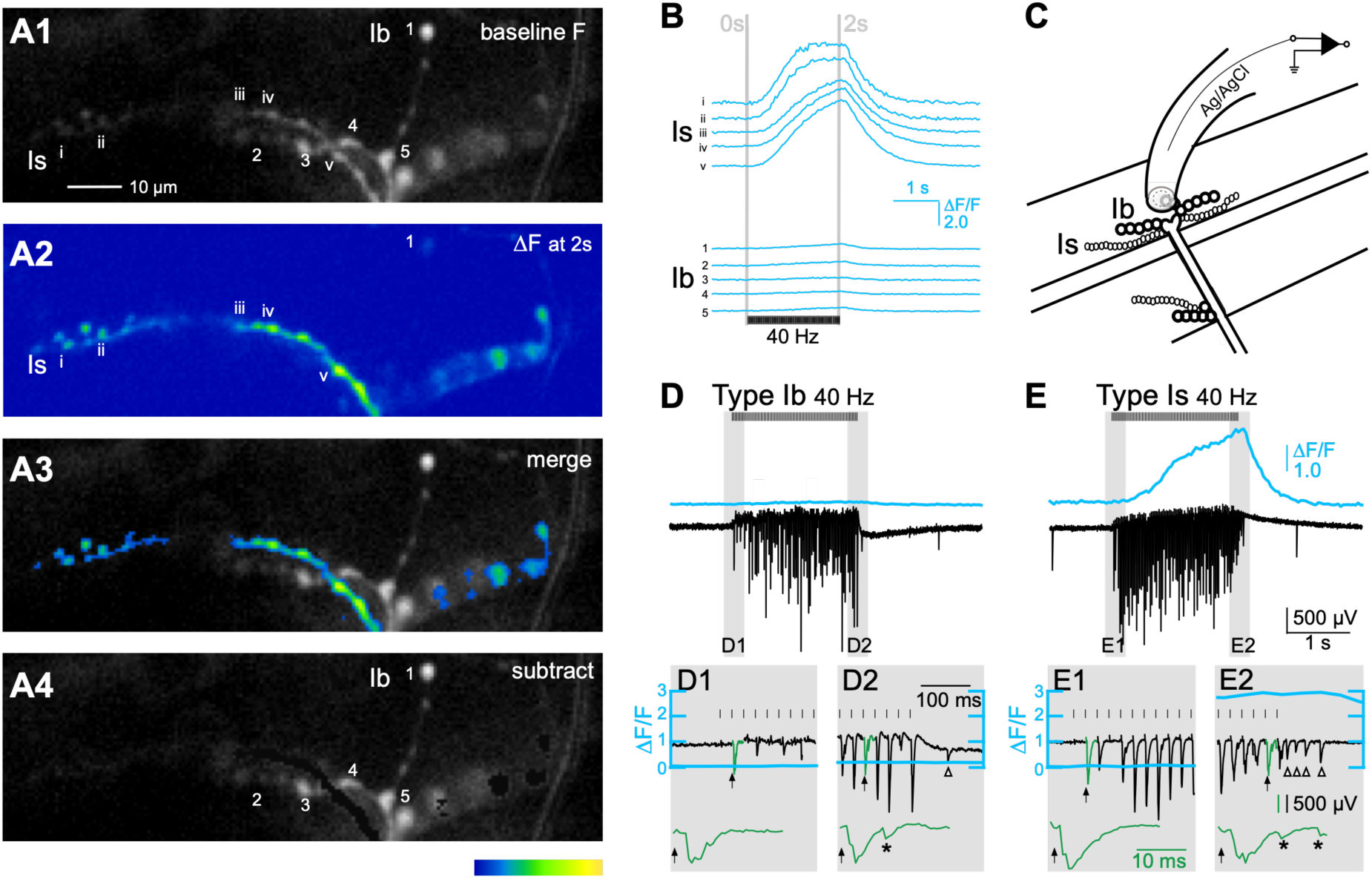
Distinct Ca^2+^ dynamics and activity-dependent plasticity in tonic Ib and phasic Is synapses in low-Ca^2+^ saline. A 40-Hz, 2-s stimulus train were delivered to nerve terminals of WT-background *c164-GCaMP6f* larvae (N = 12) in 0.1 mM Ca^2+^ HL3.1 saline. ***A1***, baseline fluorescence (F) image of the NMJ terminal before the stimulation, with representative type Ib and Is boutons individually indicated (1-5, i-v, respectively). ***A2***, heat maps that show the change in fluorescence (ΔF) immediately after (2 s) the stimulation. Responses mostly from type Is boutons (i-v). Only a little was seen from the brightest type Ib bouton (1). ***A3***, merge between A1 and A2. ***A4***, subtraction of A2 from A3 (Note the remaining type Ib boutons 1-5). Color calibration represents voltage measurements from CCD camera, from 0 to 100 mV, which applies to all the following figures. ***B***, the ΔF/F traces of the representative type Ib and Is boutons 1-5 and i-v. Note that the GCaMP signal waveform is substantially larger in type Is than in Ib. ***C***, the experimental paradigm for focal recording on NMJ boutons. Microscopic lens approaches the preparation from above (not shown). ***D***, ***E***, Focal recording of efEJP traces (black) with corresponding GCaMP ΔF/F signals (blue) simultaneously collected from the same nerve terminals (D, Tonic type Ib. E, Phasic type Is). The shaded regions (1 and 2 in each panel) are expanded in the insets (D1 and D2, E1 and E2). Note that tonic Ib showed clear enhancement (D2 vs D1) whereas phasic Is showed slight depression (E2 vs E1). Also note the supernumerary releases during (asterisks, D2, E2) and after (arrow heads, D2, E2) the stimulation train. In this and the following Figures 3, 4, 8, 10, the elevation of ΔF/F traces (blue) in the expanded panels (grey) preserves the same vertical amplitude in the original unexpanded traces. Representative efEJPs (green) are also expanded to reveal the synchronicity change.

### Simultaneous GCaMP imaging and electrophysiological recording

All solutions, equipment, protocols and data analysis methods have been described in (Xing and Wu, 2018a). Briefly, 3^rd^ instar *Drosophila* larvae were dissected in HL3 solution (Stewart et al., 1994), and recorded and imaged in HL3.1 solution (Feng et al., 2004). All data presented were collected from muscles 12 or 13, with a smaller number of samples from muscle 4 of abdominal segments 3-5. The larval segmental nerve bundles were severed from the ventral ganglion and stimulated using a glass suction electrode (7 – 10 µm inner diameter, filled with HL3.1), which contained a connecting Ag/AgCl wire. GCaMP-expressing synaptic terminals were visualized under an upright fluorescent microscope (Eclipse E600FN; Nikon) equipped with a 40X water-immersion objective lens (Fluoro; N.A. 0.80), and a high-pressure xenon short-arc lamp (UXL-75XE, Ushio). The images were collected through a CCD camera by SciMeasure Analytical Systems, and the extracellular focal excitatory junction potential (efEJP) signals were picked up through a second glass micropipette (containing an Ag/AgCl wire electrode and filled with HL3.1, see below for details) and an AC amplifier (GRASS model p15, Warwick, RI), with cut-off frequencies set to 0.1 Hz and 50 kHz. To accommodate the limited working distance beneath the water-immersion objective lens, a bent glass micropipette was used to cover and record from mostly one or occasionally 2 – 3 adjacent boutons, as visualized by the GCaMP basal fluorescence throughout the motor synaptic terminals. The electric and optic signals were integrated by RedshirtImaging NEUROCCD-SM256 system (Decatur, GA).

### Focal recording and local manipulation of synaptic bouton microenvironment

Glass micropipettes described above were pulled using a vertical puller (Model PP-83, Narishige, Tokyo, Japan), and were further heated and polished with a microforge (Series A NO 310, Microforge De Fonbrune, Aloe Scientific, MO) to have the desired openings (7 – 10 µm). For recording, the tip portion of the glass micropipettes were further heated and bent into 45 – 60 degrees so that they could easily land on muscle surface (Figure 1C).

To control the local microenvironment of the focally-recorded synaptic boutons, drugs or different ions of specified concentrations were applied to the sealed space from inside the recording pipette. This arrangement could optimize the confinement of high Ca^2+^ (or Sr^2+^) exposure to the targeted boutons and minimize muscle fiber contraction. For most of described below, higher concentration Ca^2+^ (e.g. 0.5 or 1.5 mM Ca^2+^, in HL3.1, Feng et al., 2004) was loaded in the recording pipette, whereas the bath solution (HL3.1) only contained 0 mM Ca^2+^ (Figures 3, 4) or 0.1 mM Ca^2+^ (Figure 7). Similarly, Ca^2+^ ions can be replaced with Sr^2+^ at the designate concentrations (Figures 8, 10).

All data analysis and figure construction were performed with custom-written codes based on Python. In brief, the ΔF/F measurement protocol follows that has been described in previous publications (Xing and Wu, 2018a, b). ΔF/F_2s was measured as the mean ΔF/F value between 1.95 and 2.00 s after the onset of stimulation. The heat maps of fluorescent images were converted from voltage readings of the CCD camera. No saturation of the CCD camera readout was ever encountered during the GCaMP fluorescence imaging. The size of efEJPs were measured by voltage deflections using the RedshirtImaging NEUROCCD-SM256 system. The facilitation and depression indices are described in the relevant sections in Results.

## Results

### Simultaneous GCaMP imaging and focal recording reveals distinctions in Ca^2+^ dynamics and activity-dependent plasticity between tonic Ib and phasic Is synapses

The two types of glutamatergic excitatory synaptic boutons, tonic type Ib (“I big”) and phasic type Is (“I small”), originating from different motor neurons and displaying identifiable morphologies, can be readily differentiated using Nomarski optics or fluorescence markers (Johansen et al., 1989; Atwood et al., 1993; Hoang and Chiba, 2001). Upon repetitive stimulation, tonic type Ib synapses are known to exhibit more pronounced facilitation at lower Ca^2+^ levels, and phasic type Is synapses are more prone to depression at higher Ca^2+^ levels (Betz, 1970; Kurdyak et al., 1994; Lnenicka and Keshishian, 2000).

As we previously reported, at 0.1 or 0.2 mM external Ca^2+^ in bath, a repetitive stimulation protocol of 20-40 Hz is optimal for characterizing GCaMP signal dynamics for type Ib and Is synapses without introducing movement artifacts from muscle contraction (Xing and Wu, 2018a, b). Here we present simultaneous GCaMP imaging together with focal efEJP recording to correlate the synaptic transmission process with GCaMP Ca^2+^ dynamics in type Ib and Is boutons collected from muscle 12 (Figure 1). Our results show largely consistent GCaMP signal waveforms among the boutons along the same terminal branch (see for example, labeled type Ib, 1-5, or Is, i-v, boutons in Figure 1A with corresponding ΔF/F in Figure 1B). At low Ca^2+^ levels (around 0.1 mM), GCaMP signals could be collected reliably without movement artifacts even at high-frequency stimulation (20-40 Hz, Supplement Video 1). We also confirmed a minor distal-proximal gradient of ΔF/F amplitude (Figure 1B, see also; He et al., 2009; Peled and Isacoff, 2011; Xing and Wu, 2018b). In addition, our results consistently showed that phasic type Is terminals had much larger increase in GCaMP fluorescence (ΔF/F) than tonic type Ib synapses at 0.1 mM Ca^2+^ (Mean ± SDs at the maximum ΔF/F, Ib: 0.23±0.24, n = 8; Is: 3.82±1.55, n = 4, Mann-Whitney U test, p = 0.004). Furthermore, we set out to examine the possibility that GCaMP signals could yield information relevant to activity-dependent plasticity, e.g. facilitation and depression, since the ΔF/F signals display a slow, gradual rise and are not directly correlated with the instantaneous evoked transmitter release (Figure 1B, see also Xing and Wu, 2018a and b).

To achieve simultaneous efEJP recording, given the limited working distance beneath the water-immersion objective lens, we used a bent glass micropipette to record focally from one or occasionally up to 2-3 adjacent boutons, as guided by GCaMP basal fluorescence (Figure 1C).

The transmitter release, as indicated by the efEJP responses, displayed progressive increase or decrease during the stimulation train, reflecting characteristics of synaptic facilitation or depression, respectively.

In tonic type Ib synapses, 40-Hz stimulation in 0.1 mM Ca^2+^ saline could trigger an overall progressive increase in transmission amplitude along with some stochastic fluctuation, which was accompanied with only gradual but minimal increases in the GCaMP signal (Figure 1B, D, D1, D2, cf. Xing and Wu, 2018b). In contrast, phasic Is boutons exhibited an even more evident kinetic disparity between the GCaMP signal and synaptic transmission: the gradual increase in GCaMP fluorescence lagged behind transmission for hundreds of milliseconds. When GCaMP signal (ΔF/F) was still in a latent period, i.e. showing little increase above the baseline (Figure 1E1), a steep increase in efEJP amplitude occurred within tens of milliseconds following the onset of stimulation, which falls in the typical temporal ranges of facilitation (Zucker and Regehr, 2002). Following the peaking of transmission in Is boutons, the initial rise turned into a decline phase while the GCaMP signal continued to accumulate (Figure 1E, E1, E2). When the rising GCaMP signal was plateauing, the transmission also steadily declined, despite some stochasticity (Figure 1E). However, it is noteworthy that, such transition from facilitation to depression was seen only in type Is phasic, but not in type Ib tonic, synapses and occurred at an external Ca^2+^ concentration as low as 0.1 mM. Interestingly, we found that the rise of GCaMP signals was also correlated with an increase in supernumerary, asynchronous vesicular release in both type Ib and Is synapses (asterisks, Figure 1D2, E2). Especially in type Is, a period of dispersed lingering releases was often seen after the cessation of stimulation but subsided before the return of GCaMP fluorescence to the baseline (open arrow heads, Figure 1E2; see also below).

### Higher external Ca^2+^ causes muscular contraction distorting Ca^2+^ imaging and hindering stable focal recording

A common practice for GCaMP imaging the *Drosophila* larval NMJ at higher Ca^2+^ levels is to use glutamate to desensitize the postsynaptic glutamate receptor channels, in order to prevent muscle contraction-induced movement artefacts (Macleod et al., 2002; Reiff et al., 2005; Lnenicka et al, 2006). To collect focal efEJPs together with GCaMP responses, our previous reports have refrained from glutamate application but have been restricted to 0.1 mM Ca^2+^ to avoid muscle contraction (Xing and Wu, 2018a). We attempted to extend the study to higher Ca^2+^ levels but encountered signal distortions due to muscle contractions. With bath saline containing 0.2 mM Ca^2+^, we were still able to monitor a portion of preparations without significant interference from muscle contraction (Supplement Video 2). In these samples, we obtained results consistent with observations at 0.1 mM Ca^2+^, albeit larger GCaMP signals especially for type Is synapses (Figure 2A1-A4, B).

**Figure 2.**
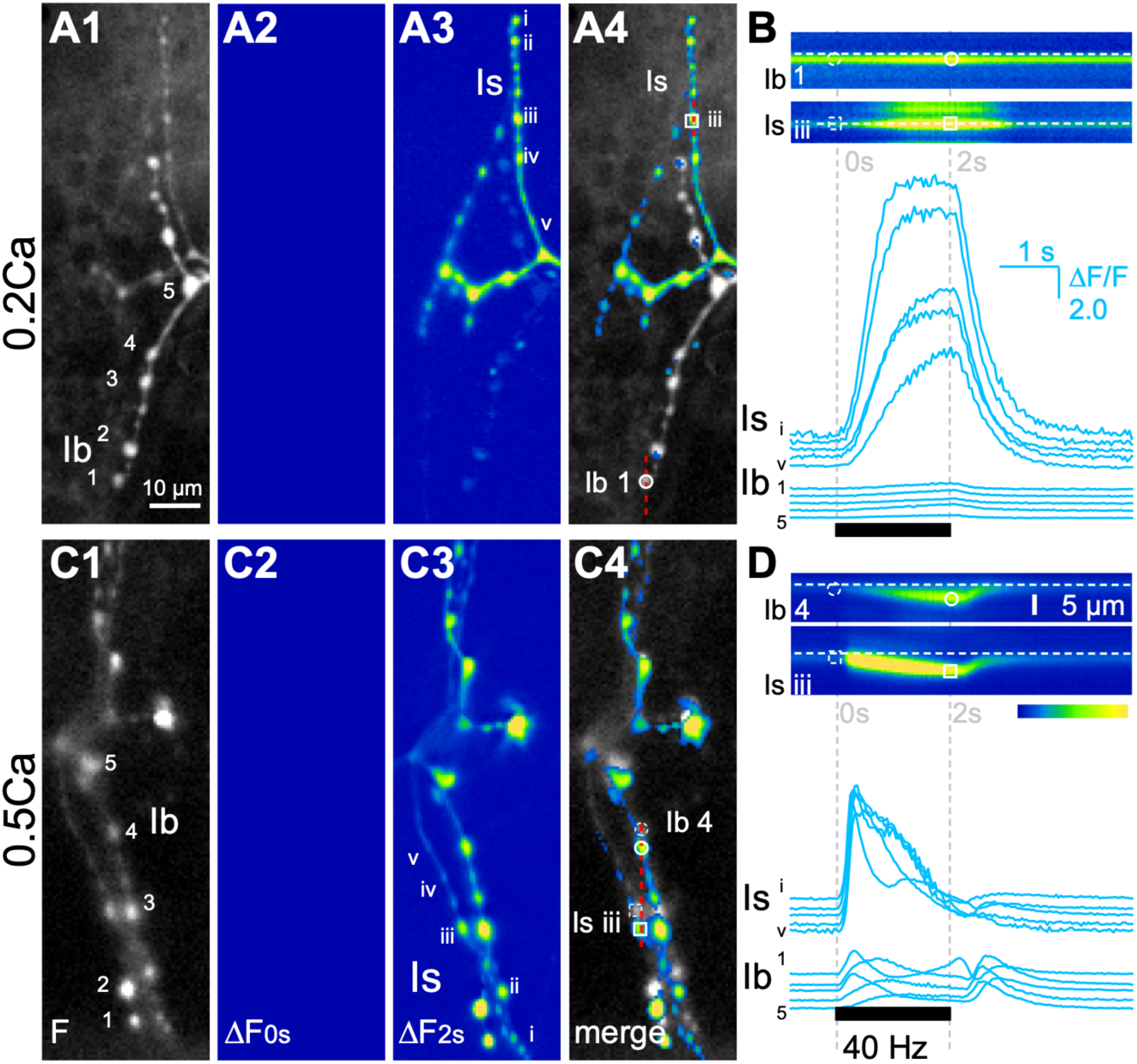
Higher external Ca^2+^ concentrations increase the occurrence of muscular contraction and distort ΔF/F signal readings. ***A1***-***A4***, Images captured from an NMJ in 0.2 mM Ca^2+^ saline, with the corresponding line-scan image (Ib 1 and Is iii) and ΔF/F signals (Ib 1-5 and Is i-v) shown in ***B***. ***C1***-***C4***, from another NMJ in 0.5 mM Ca^2+^ imaging results, corresponding line-scan image (Ib 4 and Is iii) and ΔF/F signals (Ib 1-5 and Is i-v) in ***D***. Grey vertical lines in B and D indicate the image-capture time points (0 s and 2 s) for A2 and A3, C2 and C3, respectively. Baseline F: A1 and C1; Fluorescence change ΔF is shown as heat maps at 0 s: A2 and C2, and at 2 s: A3 and C3; A4 is the merge between A1 and A3, and C4 between C1 and C3. In C4 but not in A4, note the shift in bouton position (circles: Ib, squares: Is, broken symbols: locations at 0 s, intact symbols: 2 s) due to muscle contraction. Red lines: cross-section for the line scan images in B and D. Also note the bouton movement and distortion of ΔF/F traces in D.

Without addition of glutamate to the bath, significant muscular contraction occurred at 0.5 mM extracellular Ca^2+^ (Figure 2C1-C4, Supplement Video 3). As shown in Figure 2C4, an overlay mismatches between the images before and after the stimulation (0 and 2 s), which greatly distorted the readings of ΔF/F (Figure 2D). The extent of muscular movement is indicated by the line-scan images, (derived from a cross section) tracking designated boutons along the direction of muscle movement (Ib 1and Is iii in Figure 2A4, B and Ib 4 and Is iii in Figure 2C4, D). In 0.5 mM Ca^2+^, such muscular movement (Figure 2C4, D) precludes accurate ΔF/F determination and compromises reliable focal recording from the corresponding bouton.

### Local manipulation of microenvironment extends operational range of Ca^2+^ concentrations for simultaneous GCaMP imaging and focal recording

For the purpose of stable simultaneous Ca^2+^ imaging and focal recording of synaptic transmission at higher Ca^2+^ concentrations, the protocol of glutamate application is unsuitable, which suppresses muscle contraction but also suppresses the efEJP signal (Macleod et al., 2002; Reiff et al., 2005; Lnenicka et al., 2006). Therefore, we developed a local manipulation technique, by using Ca^2+^-free bath solution and provide Ca^2+^ for transmitter release and GCaMP imaging only within the patch area via the focal recording pipette over the targeted boutons (Figure 3, cf. Figure 1C).

**Figure 3.**
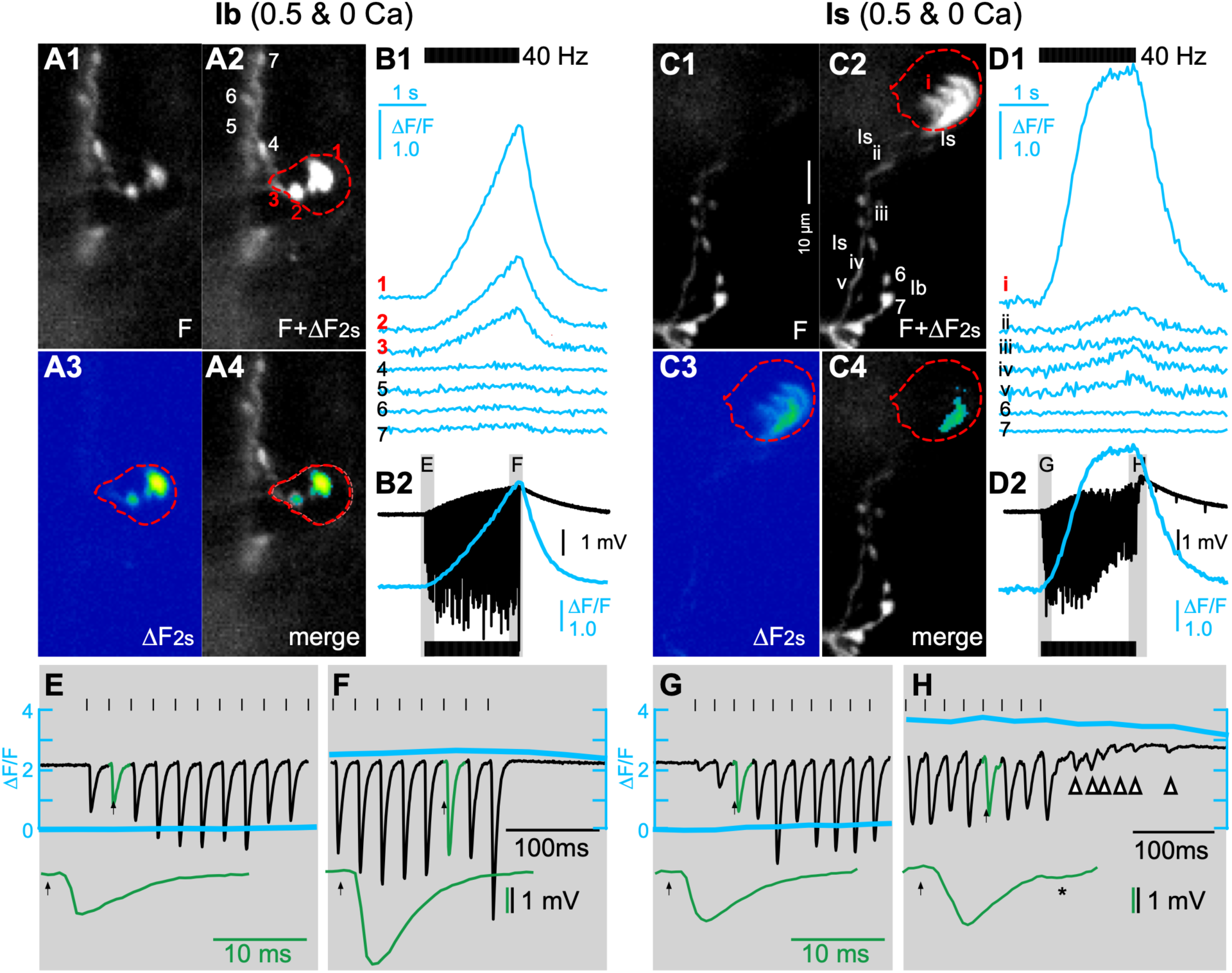
Local Ca^2+^ application in Ca^2+^-free saline enables simultaneous GCaMP imaging and focal recording of transmission without distortion by muscle contraction. Micropipette internal solution: HL3.1 with 0.5 mM Ca^2+^. Bath: HL3.1with 0 mM Ca^2+^. The pipette tip was placed either on type Ib terminal (left, ***A1***-***A4***) or type Is terminal (right, ***C1***-***C4***). ***A1***, ***C1***: baseline F before the stimulation. ***A2***, ***C2***: F at 2 s, i.e. F+ΔF_2s_, immediately after stimulation (2 s long), with representative boutons indicated (Ib 1-7, Is i-v & Ib 6, 7). ***A3***, ***C3***: ΔF at 2 s shown as heat maps. ***A4***, ***D4***: merge between the heat maps of ΔF at 2s and original F images. Note that only the boutons within the red circles (representing contours of pipette opening) showed increased fluorescence after stimulation. ***B1*** and ***D1***, ΔF/F traces measured from boutons 1-7 of type Ib (B1) and i-v of type Is and 6,7 of type Ib (D1) terminals. Red digits indicate the traces from the boutons within the pipette contours. ***B2*** and ***D2***, efEJPs measured from the encircled type Ib 1 and Is i boutons, respectively. The shaded regions are expanded together with the GCaMP response in ***E***, ***F***, ***G***, ***H***. Note the facilitation in E (tonic Ib) and G (phasic Is), and the depression (compare to G) in type Is (H). In panel H, also note the supernumerary lingering releases during (asterisk) and after (arrow heads) the stimulation train.

This arrangement successfully avoided muscle contraction in all of our experiments at 0.5-1.5 mM local Ca^2+^ (e.g. Figures 3, 4, Supplement videos 4, 5 for 0.5 mM local Ca^2+^) and yielded temporally correlated fluorescence signals and efEJPs from the individual targeted boutons (either single boutons or tight clusters of 2-3 adjacent boutons, e.g. Figures 3B1–B2, 3D1–D2, 4E, 4F). As Ca^2+^ was only applied from within the recording micropipette, the GCaMP ΔF/F signal amplitude was highest within the pipette, and declined sharply along the same terminal just outside the recording site (Figures 3A2–A4, 3B1, 3C2–C4, 3D1, Supplement videos 4, 5).

**Figure 4.**
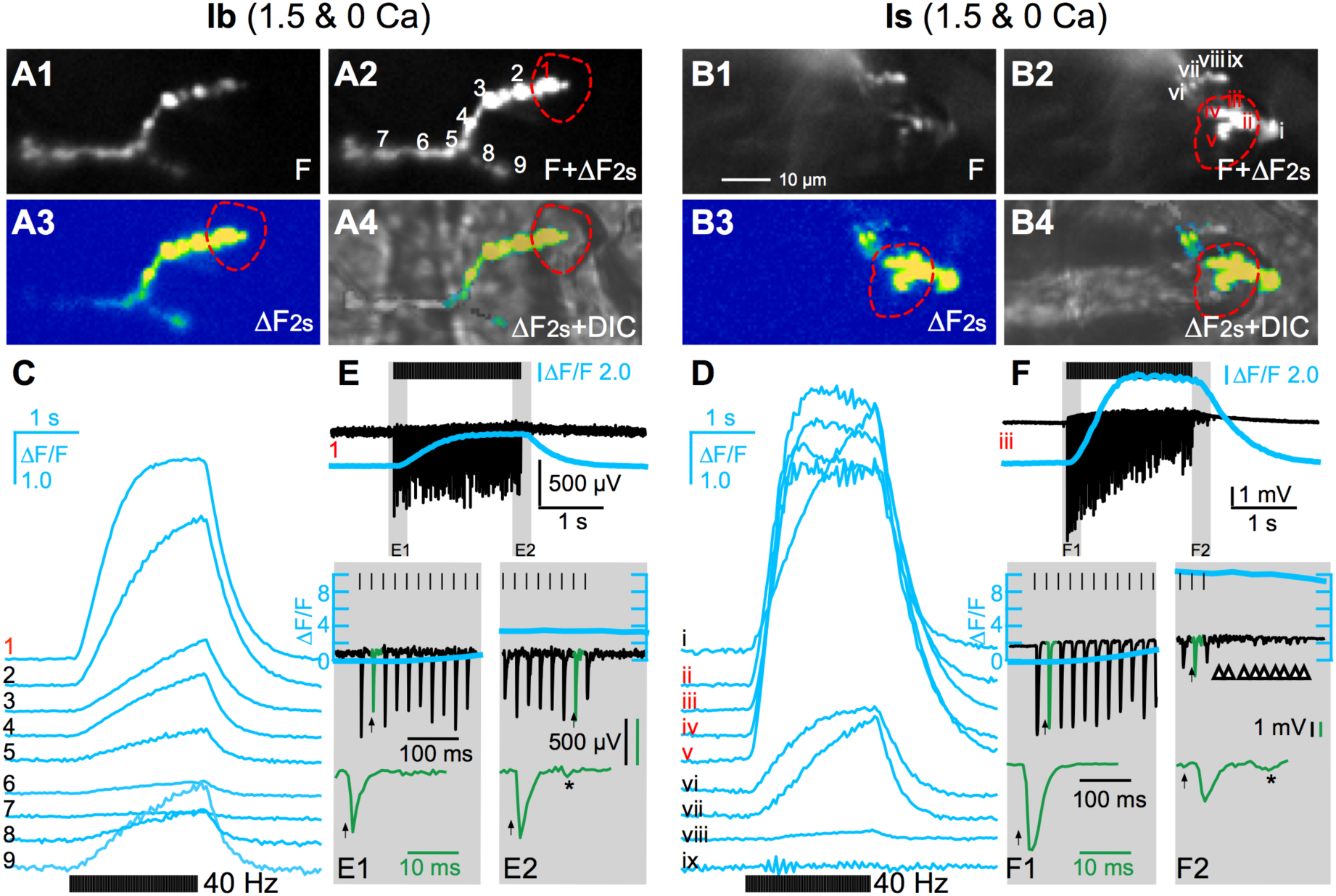
Local high Ca^2+^ abolishes facilitation and promotes depression in tonic and phasic terminals. Micropipette internal solution: 1.5 mM Ca^2+^. Bath: 0 mM Ca^2+^. The pipette tip was placed on terminals to cover the boutons in red circles (***A1***-***A4***, type Ib. ***B1***-***B4***, type Is). ***A1***, ***B1***: Baseline Fluorescence (F). ***A2***, ***B2***: F at 2 s. ***A3***, ***C3***: ΔF at 2 s shown as heat maps, color calibration same as Figure 1. ***A4***, ***B4***: merge of the ΔF images with DIC micrographs. The representative boutons are indicated as (Ib 1-9, Is i-ix). ***C*** and ***D***, ΔF/F traces measured from the indicated boutons. Red digits indicate the traces from the boutons within the pipette contours. ***E*** and ***F***, efEJPs measured from the encircled type Ib 1 and Is iii boutons, respectively, together with the corresponding ΔF/F traces from the indicated boutons. The shaded regions are expanded in ***E1***, ***E2***, ***F1***, ***F2***, with subgroups of efEJPs further expanded as green traces. Note the supernumerary releases during (asterisks) and after (arrowheads) the stimulation train in E2 and F2.

When Ca^2+^ was further increased to 1.5 mM within the pipette, we detected slightly increased spreading of GCaMP ΔF/F increments in adjacent boutons outside the pipette, possibly due to diffusion of Ca^2+^, while the muscles remained quiescent (Figure 4). Synaptic boutons at this range of Ca^2+^ concentrations showed potent levels of transmission under the micropipette opening (encircled regions, Figures 3B2, 3D2, 4E, 4F). Under this experimental condition, the GCaMP ΔF/F amplitude remained stronger in type Is than in type Ib synapses in 0.5 mM Ca^2+^ (Ib: 1.62±1.91, n = 12, Is: 4.49±1.44, n = 7, Mann-Whitney U test, p = 0.005) and showed further increases in amplitude, but less pronounced differences in 1.5 mM (Ib: 3.37±2.04, n = 10, Is: 6.59±7.22, n = 12, Mann-Whitney U test, p = 0.339). The amplitude and kinetics of individual efEJPs (reflecting excitatory junction currents, i.e. EJCs, cf. Xing and Wu, 2018a, b; Ueda et al., 2022) could be clearly resolved even at 40 Hz stimulation (see the expanded traces in Figures 3E–H, 4E1-E2, 4F1-F2). It is also noteworthy that even at an increased Ca^2+^ concentration (0.5 and 1.5 mM in the pipette), the onset of transmission occurred hundreds of milliseconds before any significant rise of GCaMP signals (Figures 3E, 3G, 4E1, 4F1), supporting the earlier notion that GCaMP signal mainly represents cytosolic residual Ca^2+^ accumulation (Xing and Wu, 2018a, b).

There was no clear correlation between the instantaneous amplitude of GCaMP signal and the size of concurrent efEJPs. However, it is again notable that increased chances of supernumerary releases were seen during and after the late phases of stimulation (Figures 3H, 4E2, and 4F2; cf Figures 1D2, 1E2), which were correlated with the plateaus of the GCaMP signals.

### Cytosolic Ca^2+^ dependence and temporal characteristics of short-term plasticity in tonic and phasic synapses

With the extended range of Ca^2+^ concentrations, we were able to obtain a more complete picture of the Ca^2+^ dependence and temporal characteristics of short-term plasticity for both tonic and phasic synapses. Upon examining the GCaMP and efEJP data together (Figure 5), we could divide the 40-Hz, 2-s train into two phases, an initial phase 0-0.25 s and a later phase (0.25-2 s), as separated by dashed lines in Figure 5. This division of time demonstrates a great discrepancy between the initial GCaMP signal and transmitter release from type Ib and Is synapses and preserves the saliant time point of efEJP shifting from facilitation to depression (Figure 5D, see also below Figures 7C, 7D, 9C, 9D).

**Figure 5.**
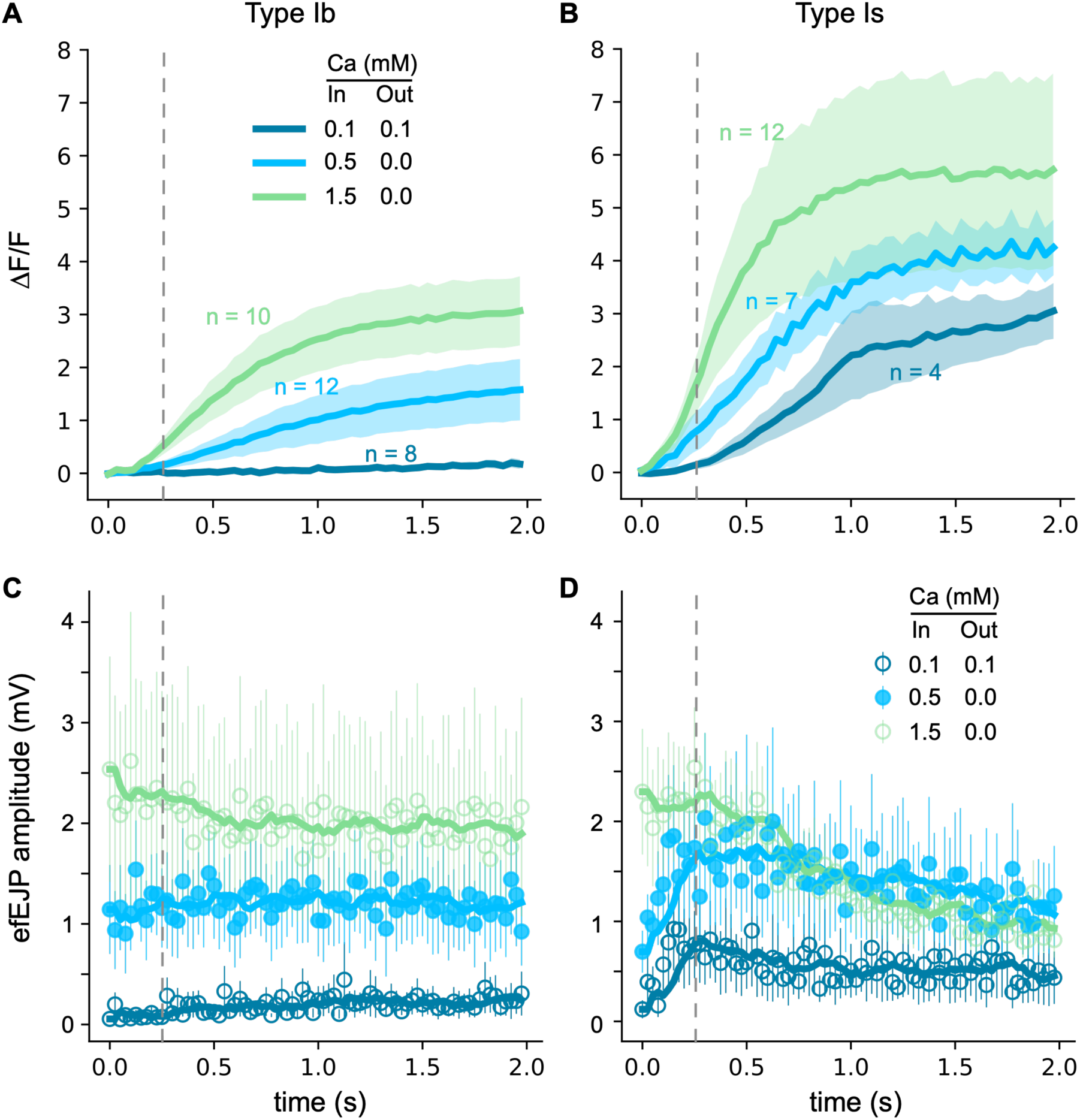
Summary plots for temporal correlation of GCaMP ΔF/F signals and efEJPs from low to high Ca^2+^ concentrations. ***A*** and ***B***, averaged ΔF/F curves for type Ib and Is boutons, respectively. For each recording, only the bouton that was under the pipette and with the strongest ΔF/F response was used for quantification. Stimulation: 40-Hz, 2-s train. Ca^2+^ concentrations as indicated. Shading represents SEM. ***C*** and ***D***, each circle represents the averaged amplitude of efEJPs recorded at the given time point. SEMs indicated as error bars. Thickened Trend lines: running average (bin = 5) of the circles. Dashed vertical lines separate the initial period (0.25 s) and the later phase. Data of each Ca^2+^ concentration group were collected from 4 – 6 larvae. Bouton numbers (n) indicated in the upper plots.

For a first-order approximation, we performed linear regression for the efEJPs during the initial and later phases separately, to obtain the mean rate of change in amplitude of the efEJPs in the two phases (in mV/s, Figure 6). The sign of the change rate indicates the tendency of facilitation (positive) or depression (negative). In 0.1 mM Ca^2+^, phasic type Is exhibited significantly stronger facilitation than tonic type Ib synapses (Figure 6A, left). Whereas at 0.5 mM Ca^2+^, the difference was weakened as type Ib showed increased tendency of facilitation (Figure 6A, middle). Increasing Ca^2+^ to 1.5 mM caused depression in both tonic and phasic synapses during the initial period (0-0.25 s, Figure 6A, right), yet phasic type Is synapses attained more severe depression than type Ib (Figure 6A, right).

**Figure 6.**
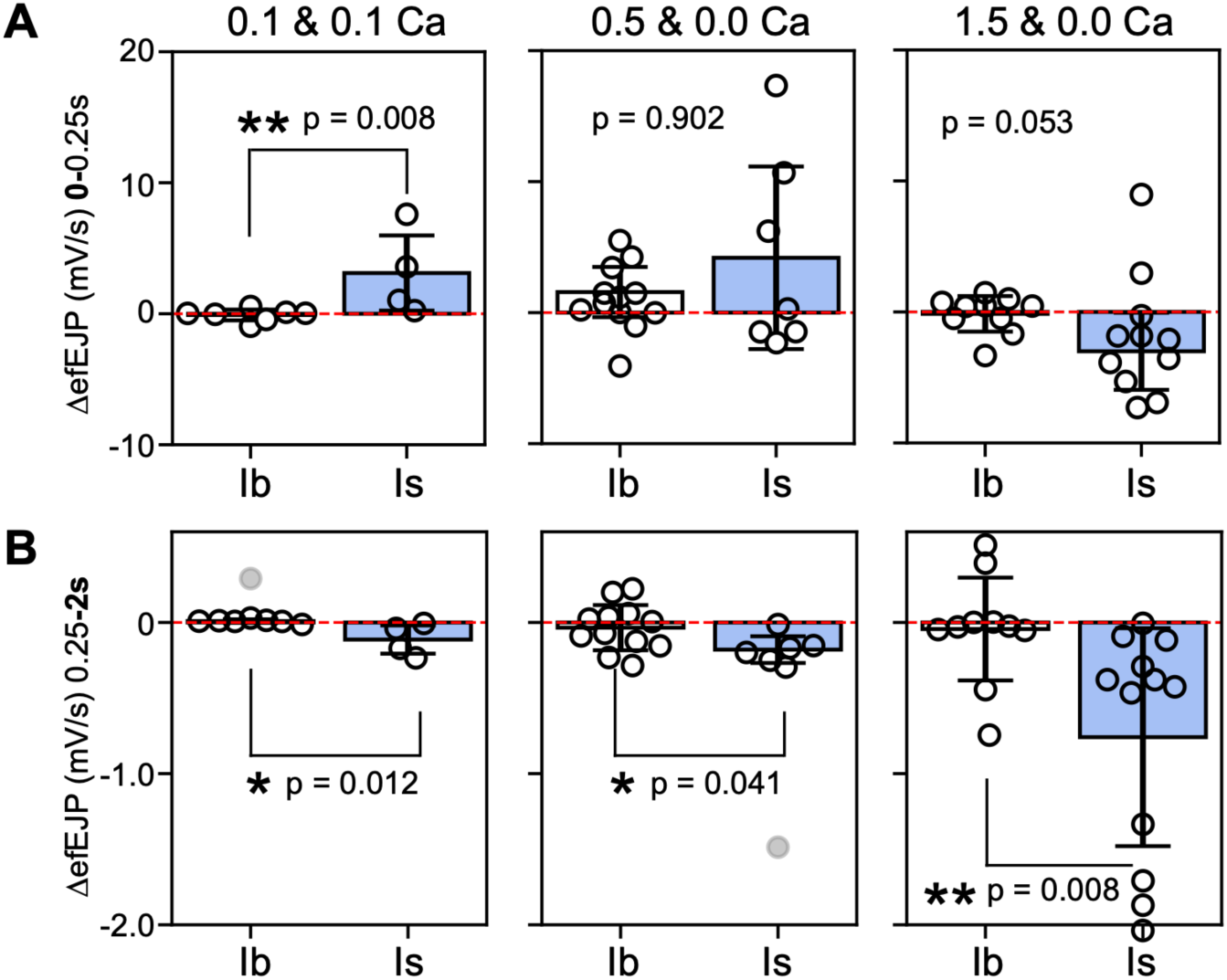
The change rates for efEJP amplitudes during the initial 0.25 s and the later phase of the 40-Hz, 2-s stimulation train. ***A***, initial (0-0.25 s). ***B***, later (0.25-2 s). Y axis: slope values obtained by linear regression of the efEJPs during the indicated time period. Each circle represents a recording from an NMJ. Positive slope: rising in amplitude. Negative slope: decreasing. Grey circles: outliers. Bars: means. Error bars: SDs. Ca^2+^ concentration in & out of the pipette as labeled above each panel. Mann-Whitney U test: *, p < 0.05; ** p < 0.01.

In the later phase (0.25-2 s), efEJP amplitude in the phasic type Is synapses exhibited stronger depression than type Ib under all conditions (Figures 6B). These observations support the view that increased residual cytosolic Ca^2+^, as indicated by GCaMP signals, would increase the release probability in the early phase (0-0.25 s) but could promote synaptic depression over facilitation in the later phase. Overall, phasic type Is synapses exhibited greater plasticity, showing steeper facilitation and stronger depression than tonic type Ib synapses (Figures 5, 6).

In contrast to 40 Hz, a lower frequency (20 Hz) of stimulation resulted in weaker expression of synaptic plasticity in both tonic and phasic synapses, exhibiting either no clear trend of efEJP amplitude change or facilitation only during the 2-s train of stimulation (Figure 7). Since 40 Hz readily induced both facilitation and depression, together with high level of GCaMP signals, this frequency of stimulation was adopted here for synaptic plasticity analysis.

**Figure 7.**
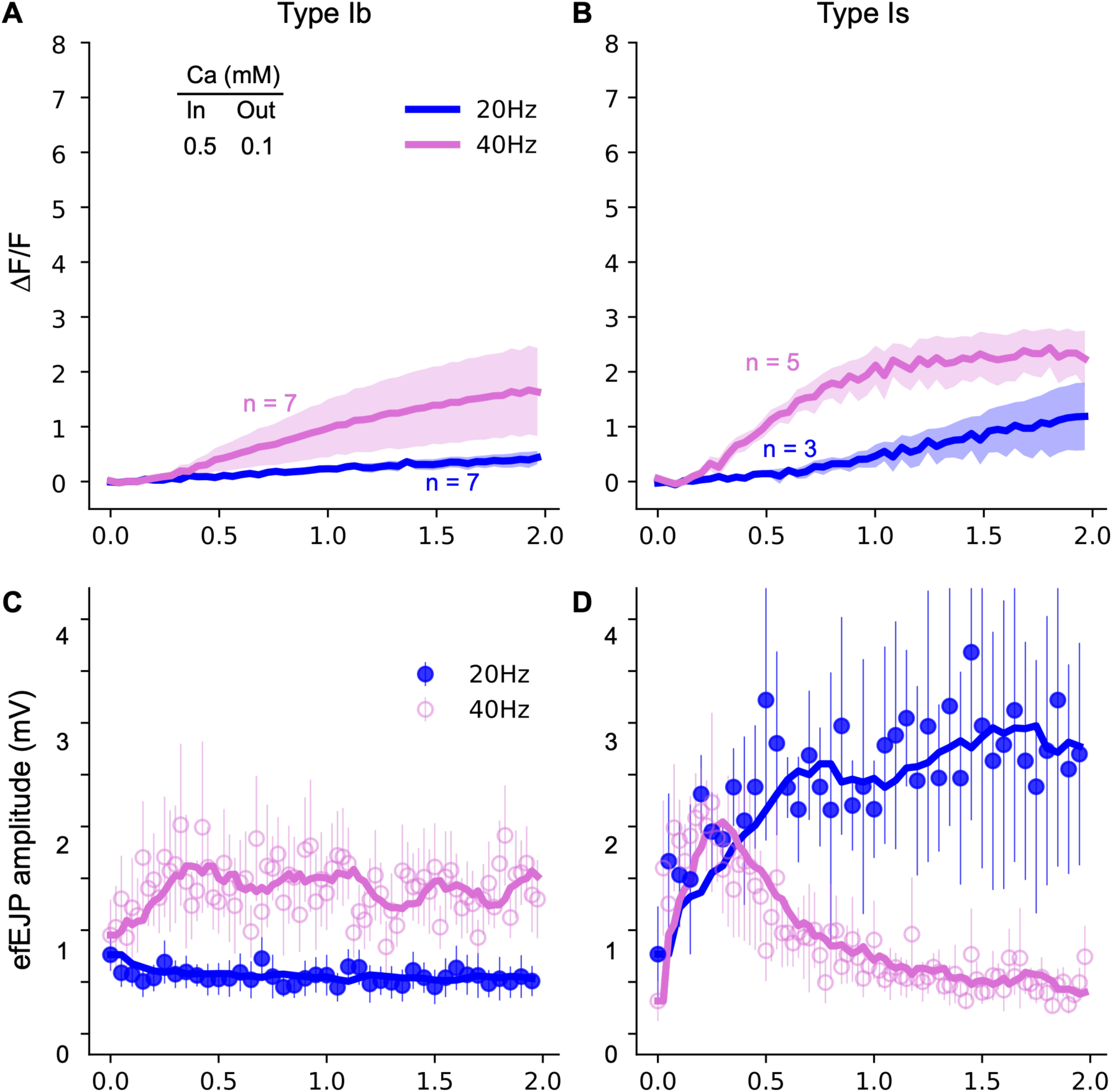
Frequency dependence for GCaMP ΔF/F signals and efEJPs of tonic type Ib and phasic Is boutons. Ca^2+^: 0.5 mM inside and 0.1 mM outside of the micropipette. Stimulation frequency as indicated. ***A*** and ***B***, averaged ΔF/F curves for tonic type Ib and phasic Is boutons, respectively. Shading represents SEM. ***C*** and ***D***, each circle represents the averaged amplitude of efEJPs recorded at the given time point. SEMs indicated as error bars. Thickened curves: running average (bin = 5) of the circles. Bouton numbers (n) indicated in the upper plots.

### Further distinctions in GCaMP signal and transmission characteristics between Ib and Is boutons revealed by replacement of Ca^2+^ with Sr^2+^

The divalent cation Sr^2+^ is well-known to pass through Ca^2+^_channels and trigger synaptic vesicular releases (Miledi, 1966; Rahamimoff and Yaari, 1973; Jan and Jan, 1976). However, to the best of our knowledge, Sr^2+^-mediated GCaMP signals have rarely been explored in NMJs (cf. Chouhan et al., 2012). To determine how Sr^2+^ retains the correspondence between GCaMP signal and synaptic transmission, we loaded the recording pipette with 1.5 mM Sr^2+^ instead of Ca^2+^, while the bath solution was devoid of Sr^2+^ and Ca^2+^. As expected, locally applied Sr^2+^ was able to effectively trigger GCaMP fluorescence increase in synapses (Figures 8). Like Ca^2+^, Sr^2+^ imaging revealed similar differences between phasic type Is and tonic type Ib synapses in terms of both Sr^2+^-induced fluorescence dynamics (maximum ΔF/F Ib: 2.76±2.17, n = 7, Is: 4.27±3.64, n = 11, Mann-Whitney U test, p = 0.328) and short-term plasticity (Figure 8). However, compared to Ca^2+^ of the same concentration, the temporal characteristics of both GCaMP signals and efEJP trains were altered in 1.5 mM Sr^2+^, showing reduced GCaMP signal amplitude, poorer transmission and thus extended period of facilitation in Ib and greater depression in Is boutons (Figure 9C, D).

**Figure 8.**
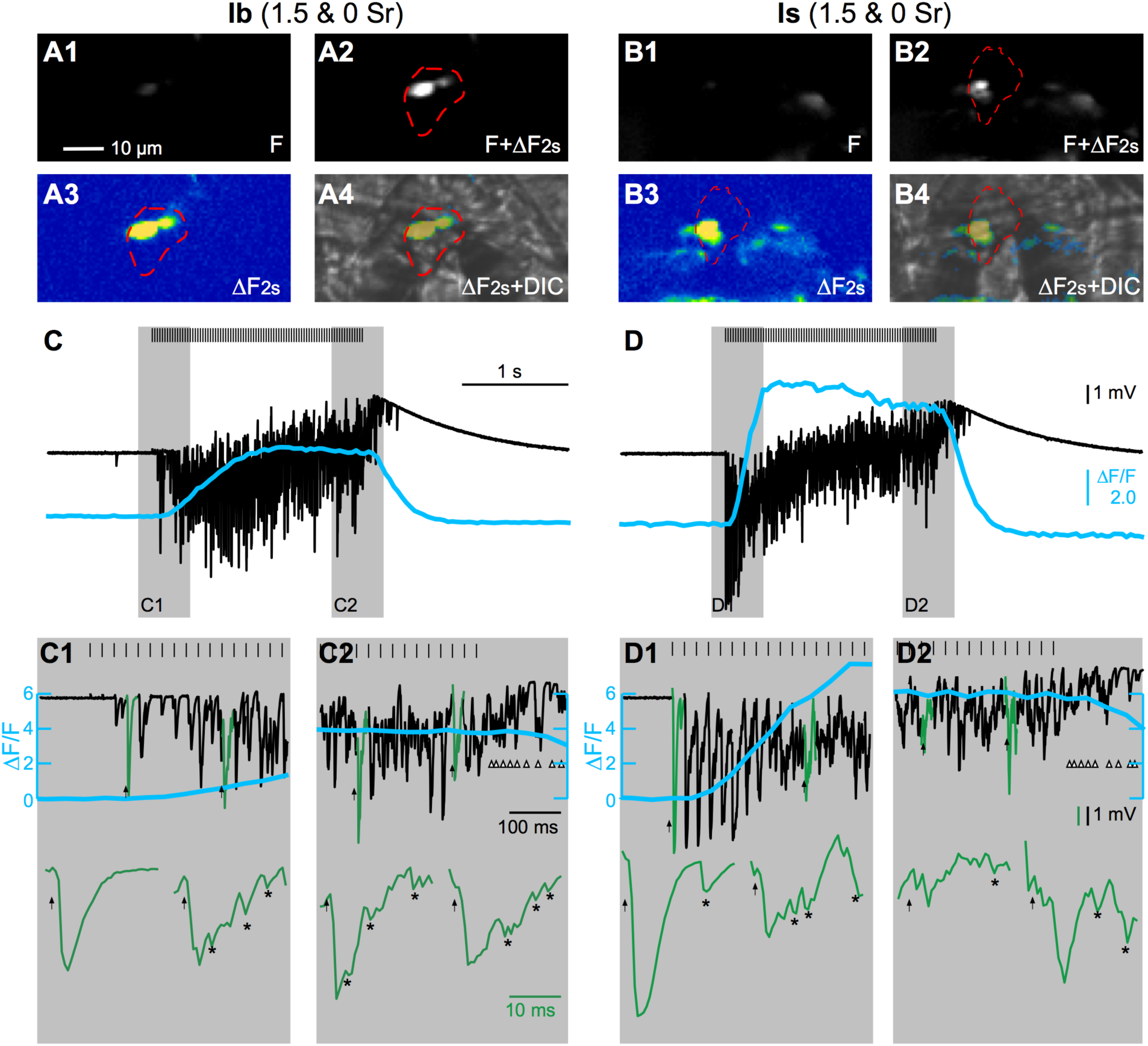
Local high Sr^2+^ (1.5 mM) differentially enhances GCaMP signals and vesicular releases in tonic and phasic synapses. Pipette internal solution: 0 mM Ca^2+^, 1.5 mM Sr^2+^. Bath: Ca^2+^- and Sr^2+^-free. 40-Hz, 2-s stimulation. Red circles indicate the pipette contours (***A1***-***A4***, type Ib. ***B1***-***B4***, type Is). ***A1***, ***B1***: baseline F. ***A2***, ***B2***: F+ΔF after the stimulation. ***A3***, ***B3***: ΔF at 2 s, subtract A1 from A2, B1 from B2, respectively, and show as heat maps. ***A4***, ***B4***: merge ΔF images with DIC images. ***C*** and ***D***, simultaneous ΔF/F signals and efEJP recordings from the encircled boutons (C: Ib, D: Is), with the shaded regions expanded as ***C1***, ***C2***, ***D1*** and ***D2***. Note the extreme asynchronicity of transmission, indicated by lots of supernumerary releases during (asterisks) and after (arrow heads) the stimulation (C1, C2, D1, D2) in both types of boutons.

**Figure 9.**
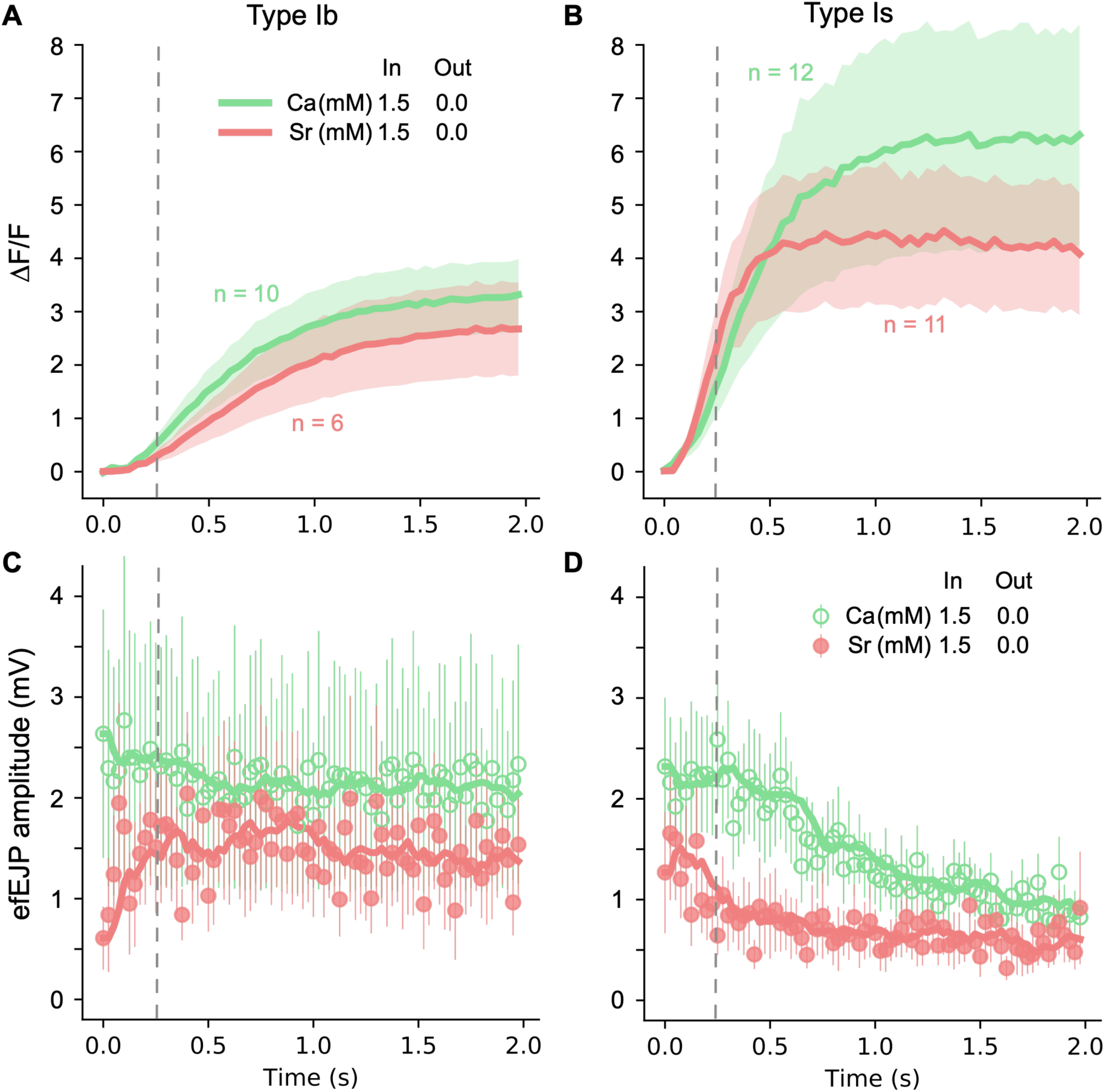
Contrasting Sr^2+^ and Ca^2+^ for induction of GCaMP ΔF/F signals and efEJPs. ***A*** and ***B***, averaged ΔF/F curves for type Ib and Is boutosn, respectively. Ca^2+^ or Sr^2+^ concentrations as indicated. Ca^2+^ data was adapted from Figure 5 for comparison. Shading represents SEM. ***C*** and ***D***, each circle represents the averaged amplitude of efEJPs recorded at the given time point. SEMs indicated as error bars. Thickened curves: running average (bin = 5) of the circles. Data of both Sr^2+^ and Ca^2+^ group were collected from 4 – 6 larvae. Bouton numbers (n) indicated in the upper plots.

Most interestingly, in Sr^2+^ solution, the rise of GCaMP Sr^2+^ signal was correlated with the onset of numerous asynchronous supernumerary releases, far more than that of Ca^2+^ (Figure 8C1, D1). During the decay phase of GCaMP signal where the stimulation has ended, a period of lingering supernumerary releases was seen to last a few hundred milliseconds and vanished before the full decay of GCaMP signals (Figure 8C2, D2). Compared to Ca^2+^ (Figures 1, 3, 4), Sr^2+^-induced post-stimulation lingering supernumerary releases were more potent and evident in both type Ib and Is synapses (Figure 8C2, D2). All these suggested that the intracellular cytosolic residual Sr^2+^, as indicated by GCaMP signals, triggered a distinct mode of vesicular release indicating the possibility of a separable vesicle pool independent of the evoked, synchronous releases (Kuromi and Kidokoro, 1998; Hagler et al., 2001; Tamura et al., 2007; Thanawala and Regehr, 2013).

Upon seeing the results of employing 1.5 mM Ca^2+^ and Sr^2+^, we sought to understand how further extended high doses of divalent cations affect the short-term plasticity of tonic and phasic synapses. However, focal application of ultra-high Ca^2+^ (3 mM) caused extensive muscular movement which prevented high-quality imaging data, even though it apparently induced severe depression even in tonic type Ib synapses. A distinct advantage of employing Sr^2+^ is that it does not effectively mediate excitation-contraction coupling (Edwards et al., 1966). We were able to obtain measurements at 10 mM of local Sr^2+^ concentration, where high Sr^2+^ completely abolished facilitation (Figure 10C1, D1), and further induced severe depression in both types of synapses Figure (10C, D). Even under drastic depression, striking supernumerary releases were also seen (Figure 10C1, C2, D1, D2). Unlike any results above, the depression was so severe that the steady-state release was reduced to a single-quantum level (Figure 10C2, D2). As a result, the short-term plasticity pattern of tonic and phasic synapses became barely distinguishable (masked by a ceiling effect). The decay phase of GCaMP Sr^2+^ signals were also extended such that long periods of lingering supernumerary releases followed the end of stimulus train (Figure 10C, D). It should be noted that at 10 mM Sr^2+^, the evoked release very rapidly goes into the depression phase without going through a massive release of transmitter (compare to Figures 3, 4, 8), suggesting that vesicle depletion could not account for the observed rapid depression, and supporting the possibility of a change in the functional state of the readily releasable vesicles.

**Figure 10.**
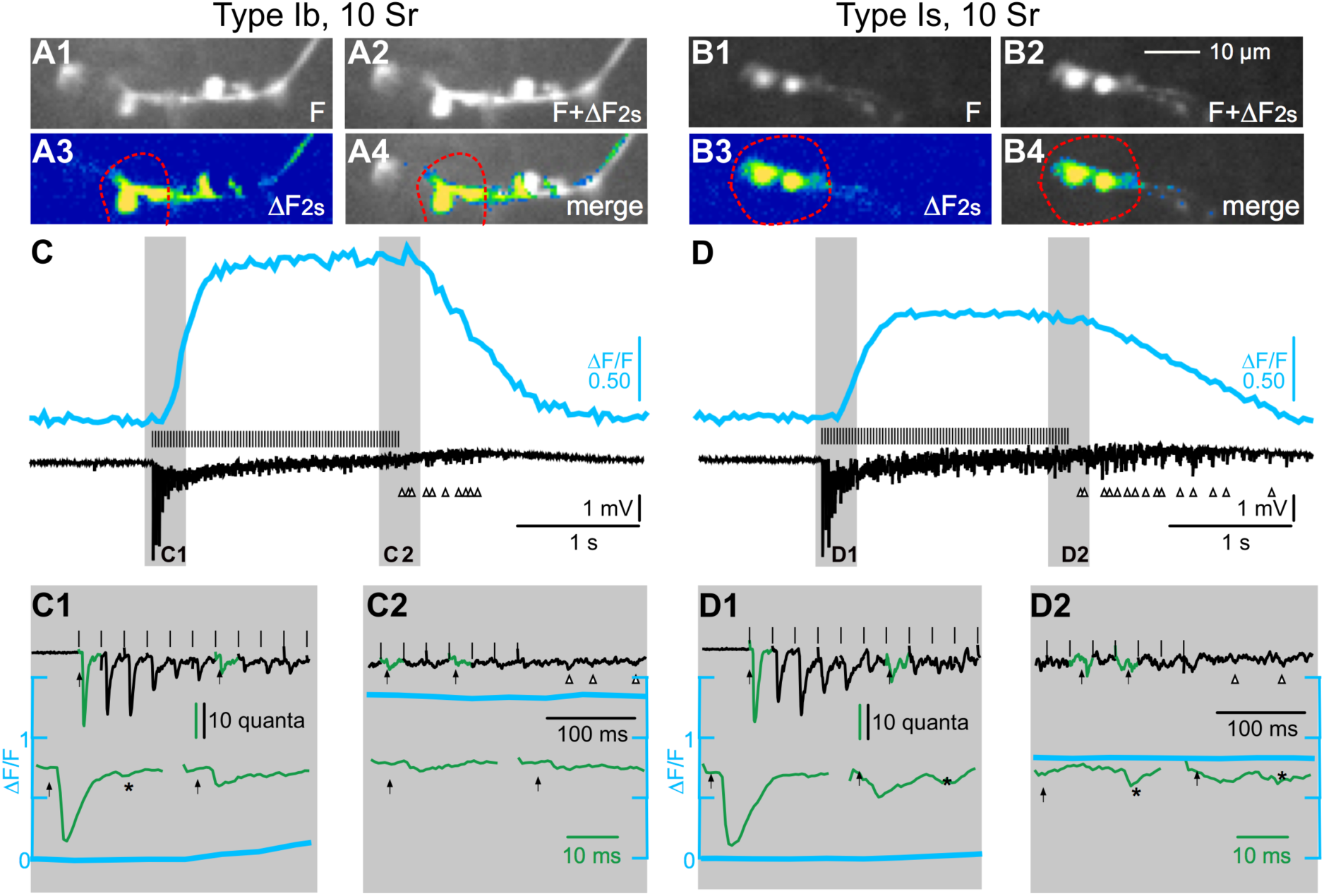
Extremely high concentration (10 mM) of local Sr^2+^ leads to elimination of facilitation and severe depression in both phasic and tonic synapses. Internal solution: 0 mM Ca^2+^, 10 mM Sr^2+^. Bath: Ca^2+^- and Sr^2+^-free. 40-Hz, 2-s, stimulation. Red circles indicate the pipette contours (***A1***-***A4***, type Ib. ***B1***-***B4***, type Is). ***A1***, ***B1***: baseline F. ***A2***, ***B2***: F + ΔF at 2s. ***A3***, ***B3***, ΔF shown as heat maps. ***A4***, ***B4***: merge ΔF images with F. ***C***, ***D***, simultaneous records of ΔF/F signals and efEJP from the encircled boutons (C: Ib, D: Is), with the shaded regions expanded as ***C1***, ***C2***, ***D1*** and ***D2***. Note the severe depression to nearly single-quantum level in both type Ib and Is synapses, and the supernumerary releases indicated by asterisks and arrow heads during and after the stimulation train, respectively.

### Global segregation of tonic and phasic boutons in Ca^2+^/Sr^2+^ dynamics and short-term plasticity characteristics

To help visualize an overall picture for distinctions between tonic Ib and phasic Is synapses, we portrayed the characteristics of GCaMP signals and the paired transmission data in a 3-D plot. As shown in Figure 11A, the X-axis represents GCaMP fluorescence ΔF/F measurements at 2s (end of the stimulation train, reflecting the net accumulation of cytosolic Ca^2+^) against the Y-axis, presenting the time to peak efEJP (or Max efEJP), which occurs either around the transition from the initial facilitation to later decline or toward the end of stimulation train for weaker efEJPs. Lastly, Z axis was used to characterize the extent or tendency of both facilitation and depression for each bouton, which was represented as a pair of indices:

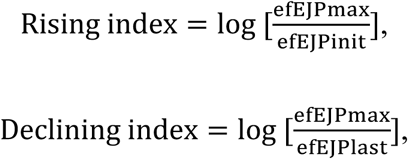

**Figure 11.**
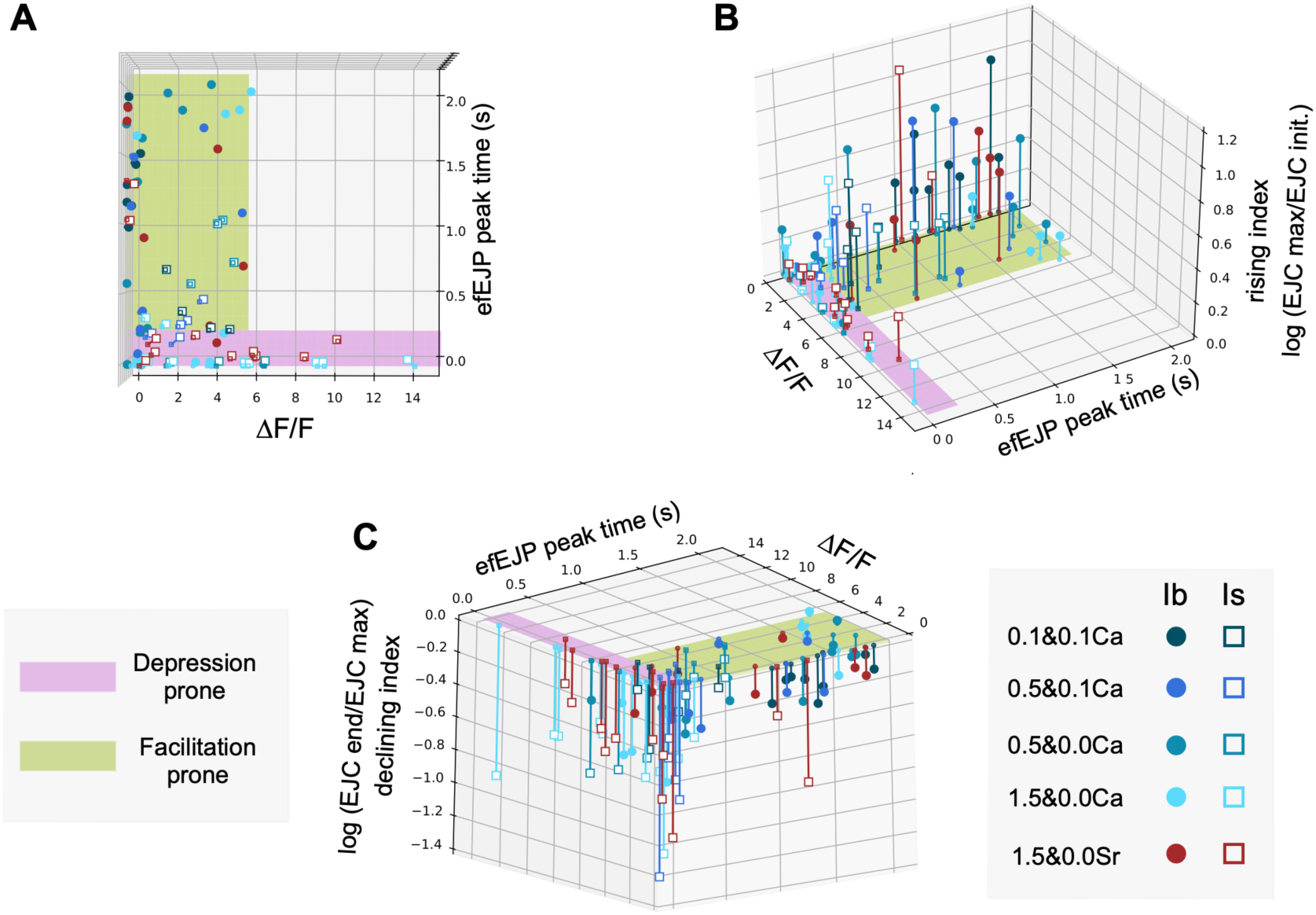
Segregation of tonic and phasic boutons in 3D plot. ***A***, Transactional view, showing the distribution of data points segregated by ΔF/F (measured at 2s after stimulation) and time points of peak efEJPs. ***B***, Positive side, efEJP rising (Facilitation) index. ***C***, Negative side, efEJP declining (Depression) index. (See text for details.) Facilitation and depression indices from the same bouton are connected by sticks, which crossed the zero plane through the mini black circles (Ib) or squares (Is). Green area (slower rising efEJPs, lower ΔF/F) contains a majority of Ib boutons that mainly exhibit facilitation. Purple area (faster rising efEJPs, stronger ΔF/F) is populated by Is boutons that mainly exhibit depression.

Where, log is log_10_, efEJP_Max_ is determined as Mean of maximum 5 consecutive efEJPs, efEJP_Init_ as Mean of initial 5 efEJPs, and efEJP_Last_ as Mean of last 5 consecutive efEJPs.

The advantage of these two logarithmic indices is that the rising index will only be positive, indicating the tendency for facilitation, or negative for a tendency of depression. Thus, the short-term plasticity characteristics of each bouton can be conveniently represented as a vertical line of the same color connecting the pair of the two points representing each bouton, with the upper showing the rising index, and the lower specifying the declining index (Figure 11B, C). If a connected pair of data points showed larger rising index but smaller declining index, the corresponding bouton was considered as “facilitation prone”, and vice versa as “depression prone”, as they appear to be separately populated in respective color-coded areas in Figure 11A.

Most tonic type Ib boutons (filled circles) showed moderate facilitation but even weaker depression, clustering in the corner with long peak EJC time and small ΔF/F amplitudes, and thus were considered as “facilitation prone” (greyish green region, Figure 11A, B, C). In contrast, most phasic type Is boutons (open squares) showed severe depression and weak facilitation, and scattered mostly in “depression prone” region (pink regions, Figure 11A, B, C).

## Discussion

The precise temporal correlation of presynaptic GCaMP signals to the stepwise sequential events in the synaptic transmission processes still await further elucidation (Peled and Isacoff, 2011; Xing and Wu, 2018a, b; Singh et al, 2018). Our data represent an initial effort to focus on the quantitative correlates between the kinetic features of presynaptic GCaMP signals and the manifestation of transmission short-term plasticity. In this report, we devised a local microenvironment manipulation protocol, which minimized interference from muscle contraction to allow for stable simultaneous optical Ca^2+^ imaging and electrical focal recording from individual synaptic boutons at a temporal resolution of milliseconds. This approach also revealed several characteristic properties in extended ranges of divalent cation concentrations for the tonic type Ib and phasic type Is synaptic terminals not previously elucidated. The translucent preparation of *Drosophila* NMJs effectively facilitates the application of this approach in further correlational investigations of GCaMP signals and related transmission activities within individual boutons. Such integrated comprehensive information can provide insight into how the activity-dependent residual Ca^2+^ accumulation modulates synaptic short-term plasticity mechanisms.

### Temporal integration of focal efEJP recording and GCaMP imaging under local microenvironment manipulation: kinetic and amplitude parameters

Our local manipulation protocol is a revamp of the classical works of focal extracellular recordings on the frog endplates by Fatt and Katz (1952) and the crayfish NMJs (Dudel and Kuffler, 1961; Atwood, 1967). A confined local microenvironment could be further created by loading the focal recording micropipette with solutions of defined compositions so that the membrane patch underneath is exposed to a designated local microenvironment but isolated from the outside bath solution. Importantly, since insect muscle contraction requires Ca^2+^ influx, this arrangement minimizes the interference from muscle movements without relying on the commonly adopted practice of inducing postsynaptic receptor desensitization by glutamate (Macleod et al., 2002; Millar et al., 2005; Reiff et al., 2005; Lnenicka et al., 2006; Melom et al., 2013), which precludes the correlation of GCaMP signals with postsynaptic responses.

Compared with intracellular recording, the focal recording directly registers the local extracellular field potentials (efEJPs) generated by the excitatory junction current flow around the targeted synaptic boutons. Thus, efEJPs have better temporal resolution than EJPs for resolving the time course of immediate postsynaptic currents, without involving the membrane capacitance element that shapes EJPs. The simple efEJP signals hence can faithfully monitor the dynamics of vesicle releases and postsynaptic responses with optimized temporal resolution, especially for the dispersion due to asynchronous releases (e.g. Figures 1D2, 1E2, 3H, 4E2, 4F2). By confining the divalent ions availability to the targeted regions under the focal pipette, this protocol enables local control of Ca^2+^ levels and, in principle, it can be modified for local drug delivery to enhance spatial and temporal correlation between GCaMP signals and drug actions within the same bouton(s). Another advantage of this simple operational protocol and experimental configuration is to facilitate rapid data collection from multiple synaptic loci along the synaptic terminal branches for fine mapping and other functional analyses.

It should be noted that this system does not have the voltage-clamp functionality to control the cross-membrane driving force, unlike focal loose-patch clamp recording (Fenwick et al., 1982; Kurdyak et al., 1994; Renger et al., 2000). Therefore, variations in pipette seal resistance and in transmembrane potential could lead to variable and uncalibrated efEJP sizes from patch to patch. Nevertheless, the absolute time course and relative efEJP size can still be monitored with precision for each patch, which faithfully reflects the plasticity pattern of the synaptic transmission (e.g. Figures 1, 3, 4, 5, see also Dudel and Kuffler, 1961; Atwood, 1967). Specifically, the spontaneous quantal events can be captured during the same recording session to serve as an internal scaling factor, so that the quantal content of the evoked releases can be estimated for each patch (Figure 10).

Absolute quantification of transmission amplitude will be best performed with the postsynaptic muscle membrane voltage-clamped (Martin, 1955; Wu and Haugland, 1985; Singh and Wu, 1990; Lee et al., 2008), because depolarization during EJPs (produced by the ensemble of all synaptic boutons on the muscle cell) reduces the driving force for ion flux through the receptor channels, especially at higher external Ca^2+^ concentrations. It has been shown such whole-cell depolarization can reach an extent to cause a considerable outward positive current component in loose patch-clamp recordings (Msghina et al., 1998; 1999). Actually, the local manipulation protocol can reduce these undesirable effects by confining the transmission current to underneath the micro-pipette, a small portion of the muscle cell membrane.

### The kinetics of GCaMP signal and its temporal correlation with short-term synaptic plasticity

The paired-pulse stimulus protocol has been regularly used in short-term plasticity studies, whereas motor neurons typically fire in trains of action potentials (Levine and Wyman, 1973; Ikeda and Kaplan, 1970; Marder and Calabrese, 1996; Fox et al., 2006). Longer trains of repetitive stimuli, which effectively activate GCaMP signals, have been widely adopted in GCaMP imaging for investigation of intracellular free Ca^2+^ levels (Reiff et al., 2005; Mao et al., 2008; Chouhan et al., 2010; Chen et al., 2013; Martin et al., 2016; Xing and Wu, 2018a, b). Our data indicate that the temporal characteristics of synaptic short-term plasticity exhibit certain correspondence in parallel with the time course of presynaptic GCaMP signals.

It should be noted that GCaMP signals described here represent a spatially averaged, global level of cytosolic Ca^2+^ within synaptic boutons and does not correspond to the near-active zone Ca^2+^ influx critical for vesicular transmitter release activity. Therefore, interpretation of these GCaMP signals may be aided by investigating their temporal correspondence to elements in the sequence of synaptic transmission and plasticity events.

GCaMP signals typically exhibit a lag of about 100-200 ms behind transmitter release, most conspicuous at low Ca^2+^ levels, with shorter lags observed at higher Ca^2+^ concentrations. Synthetic Ca^2+^ indicators, although faster than GCaMP, also exhibit some lag behind transmission (Samigullin et al., 2015). Several factors, including the kinetics of GCaMP fluorescence emission which depends on its calmodulin-binding of four Ca^2+^ ions for activation, may contribute to this lag. Such consequence is more exaggerated at low Ca^2+^ levels, which is akin to the activation of delayed rectifier K^+^ current (I_K_) evoked in response to a voltage jump across the cell membrane.

This dynamic feature is well described by the Hodgkin-Huxley (1952) model based on the assumption of four identical independent control entities (or channel subunits). Similarly, the delay in I_K_ rise is longer at smaller voltage jumps and progressively shortened when the depolarizing step increases.

Clearly, the duration of the lag phase can be greatly influenced by additional factors, such as the regulation of Ca^2+^ influx during repetitive neuronal firing. Increased extracellular Ca^2+^ concentrations undoubtedly enhance Ca^2+^ influx, thereby shortening the lag phase (Figure 5). Increased neuronal excitability can lead to stronger and more prolonged membrane depolarization, allowing voltage-gated Ca^2+^ channels to open longer and admit a greater flow of Ca^2+^. For example, blockade or mutational alteration of K^+^ channels has been shown to strongly boost Ca^2+^ influx and hasten the rise of GCaMP signals at the larval NMJs (Ueda and Wu, 2006; Xing and Wu, 2018a). It has been reported that after removal of K^+^ channel repolarization by multiple channel blockers or mutations, a single stimulus of the motor axon can induce max GCaMP response, reaching a height comparable to the plateau level in untreated larval NMJs (Xing and Wu, 2018a). This demonstrates the capability of the CaV channels in the synaptic terminal in supporting prolonged Ca^2+^ influx to elicit faster build-up of intracellular free Ca^2+^ (Katz and Miledi 1967; Ganetzky and Wu, 1982, 1983), leading to full-sized GCaMP response. However, it should be noted that even under such conditions, a lag of about 100 ms still persists in the GCaMP signal rise behind synaptic transmission, as indicated by efEJP detection (Xing, 2014; Xing and Wu, 2018a).

It is also desirable to extract a first-order estimate about the cytosolic Ca^2+^ concentrations at which these plasticity events occur. For the effective operational range of Ca^2+^ imaging, the Hill coefficient of GCaMP6m is somewhat less than 4 (2.96, cf. Chen et al., 2013), suggesting interactions among the binding sites, and resulting a steep, non-linear Ca^2+^ concentration dependence. The sigmoidal relationship shows a narrowed dynamic range (less than 2 log units) with sublinear, suppressed fluorescence responses at lower Ca^2+^ concentrations below K_d_ (Rose et al., 2014). Further, as the reported K_d_ of GCaMP6m is 167 nM (Chen et al., 2013), it can be deduced that cytosolic Ca^2+^ indicated effectively by GCaMP fluorescence is in the range of hundreds of nM. An intracellular Ca^2+^ concentration in this range is unable to trigger massive, synchronous releases of vesicles from the releasable pool, but could be adequate for triggering other cellular processes, such as facilitation or depression.

Near the active zones, rapid Ca^2+^ influx causing exocytosis within fractions of milliseconds (Katz and Miledi, 1965), a rapid event not captured by GCaMP imaging. The local Ca^2+^ clearance mechanisms, including plasma membrane Ca^2+^ ATPases (PMCA), are effective in extrusion of Ca^2+^ to terminate the transmitter release (Lnenicka et al., 2006; Roome and Empson, 2013). However, at lower external Ca^2+^ concentrations, the ensuing small rise of the residual Ca^2+^ can impose a significant increase in the efficiency of subsequent axonal action potentials in triggering release, producing larger EJPs, owing to the 4^th^ power Ca^2+^-dependence of transmitter release (Dodge and Rahamimoff, 1967). This synaptic facilitation process can occur at a sub-K_d_ Ca^2+^ concentration of GCaMP (see above) and is consistent with our observation that facilitation of efEJPs could be seen in the latent phase, before the detectable rise of the GCaMP signal (Figures 1, 3).

Under the condition of repetitive stimulation, the releasable vesicle pool must be replenished, and when the replenishment rate is insufficient, the equilibrium is leaning toward the depletion of the releasable vesicles, entering the process of depression. The transition from facilitation to depression was observed around 250 ms from the onset of the 40-Hz stimulation train, during the rising phase of the GCaMP signal (Figures 5, 7). With high external Ca^2+^, the large influxes quickly deplete the readily releasable pool of vesicles, leading to immediate depression and occasionally bypassing facilitation entirely (Figures 4, 5).

The subsequent plateau phase of GCaMP signals suggests elevated levels of cytosolic residual Ca^2+^ have reached a steady state, where the net free Ca^2+^ is balanced between influx and clearance from the cytosolic compartment, likely within the same order of magnitude or higher than the K_d_ for Ca^2+^-GCaMP association (hundreds of nanomolar). The plateau phase of GCaMP signals often aligns with the steady-state levels of efEJPs (Figure 5). High levels of GCaMP fluorescence indicate effective activation of its calmodulin component, also implying other critical molecular events, such as calmodulin-dependent inactivation of Ca^2+^ channels (Lee et al., 1999) and triggering cAMP production by adenylyl cyclase (Livingstone et al., 1984; Yovell et al., 1992). These processes are crucial for vesicular recycling and replenishment, supporting sustained synaptic transmission (Zhong and Wu 1991; Kuromi and Kidokoro, 1998). Impairment in Ca^2+^ clearance mechanisms has been shown to markedly prolong the decay phase of GCaMP signals, elevating intracellular Ca^2+^ levels, and potentially facilitating asynchronous neurotransmitter release (see below).

### GCaMP signal and transmission synchronicity

Our results at higher external Ca^2+^ concentrations revealed consistent induction of asynchronous transmission during or after the repetitive stimulation, more pronounced in type Is than Ib synapses. It either appeared as dispersed efEJPs in response to individual stimulus pulses (see green traces in Figures 1D2, 1E2, 3H, 4F2, 8C2, 8D2), or as multiple linger releases after the 2-sec stimulus train (Figures 1D2, 1E2, 3H, 4F2, 8C2, 8D2). Either way, such asynchronous transmission occurred most frequently during high levels of GCaMP signals, either toward the end or right after the cessation of the 2-s stimulation train (when cytosolic Ca^2+^ accumulation reaching or declining from a plateau). In contrast, such dispersion was rarely seen in the beginning phase of the stimulation trains (Figures 1D1, 1E1, 3E, 3G, 4E1, 4F1).

Synchronous transmission can be triggered by local Ca^2+^ influx in the vicinity of the active zones (AZ) through orchestrated opening of voltage-gated Ca^2+^ channels that are driven by membrane depolarization upon arrival of motor axon action potentials (Zucker, 1996, see also a summary diagram Figure 12).

**Figure 12.**
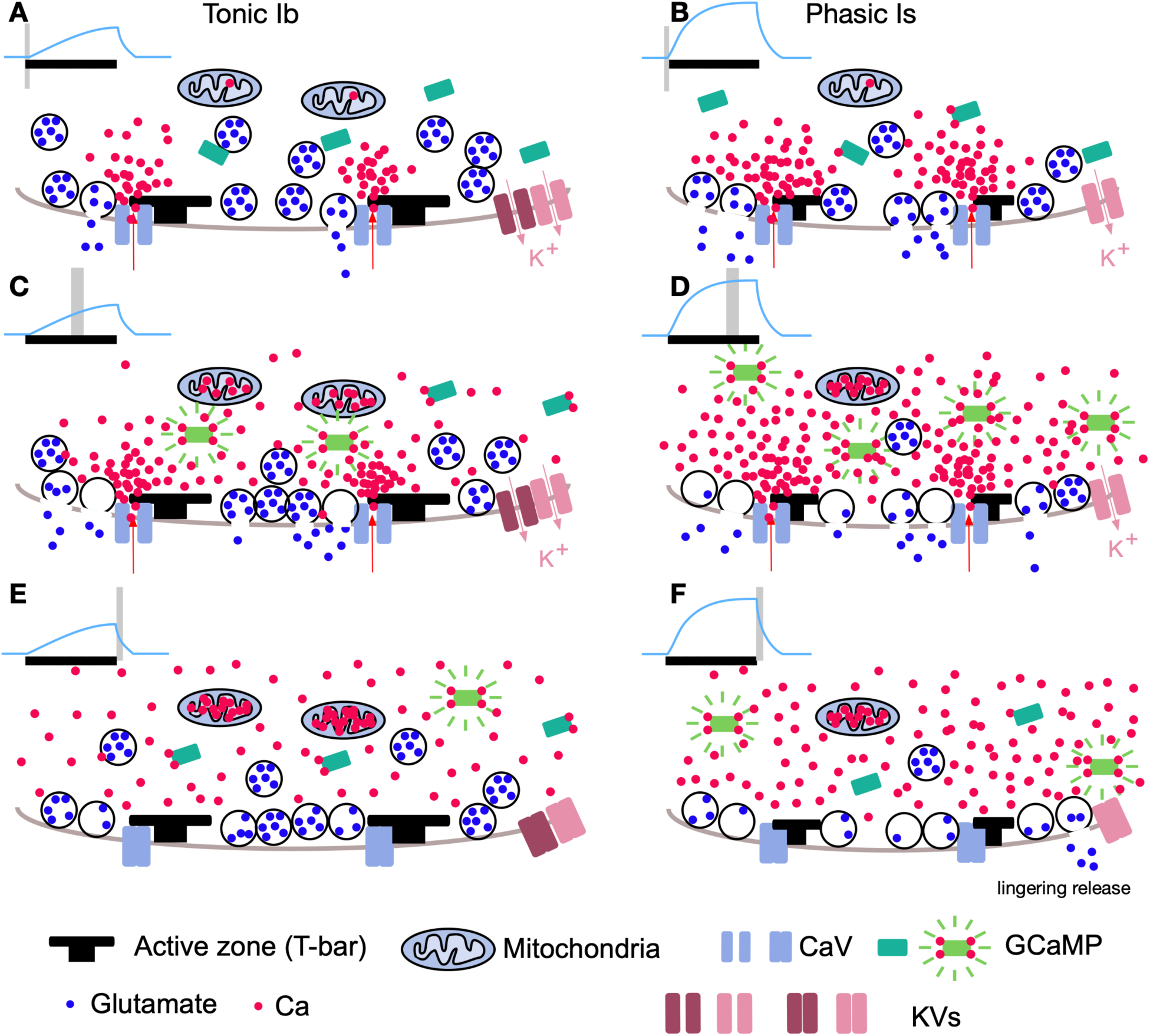
Summary for the significance of GCaMP fluorescence during the transmission aroused by repetitive stimulation. Left: illustration for tonic type Ib synapses. Right: phasic type Is synapses. ***A*** and ***B***, synchronous transmission triggered by instantaneous opening of Ca^2+^ channels upon arrival of action potentials. ***C*** and ***D***, repetitive stimulation causes accumulation of residual Ca^2+^, recruitment of more vesicular release, and depletion of releasable vesicles. The transmission may become asynchronous, i.e. dispersed, due to the recruitment of additional releasing sites. ***E*** and ***F***, after the end of repetitive stimulation, the residual Ca^2+^ may sporadically induce some vesicular exocytosis, i.e. lingering release. Note the tighter control of the vesicular release and residual Ca^2+^ clearance processes in Ib boutons, owing to a higher density of mitochondria, a greater K^+^ channel number or variety, and potentially different ultrastructure of the T-bar (active zone).

As mentioned above, the effective cytosolic Ca^2+^ levels reported by GCaMP fluorescence is in the range of hundreds of nM. An intracellular Ca^2+^ concentration in this range is well below the levels (transiently greater than 10 µM, Zucker, 1996; Sun et al., 2024) to trigger massive, synchronous releases of vesicles from the releasable pool, but might be adequate for triggering a small portion of synaptic vesicles to be released sporadically upon regional upward fluctuation of local Ca^2+^ levels (diagramed in Figure 12F). It should be stressed that this may occur not only near the AZ but also in regions farther away, as different modes of vesicular release at sites away from the AZ have been suggested previously for spontaneous or asynchronous release (Sakaba and Neher, 2001; Tamura et al, 2007; Melom et al., 2013; Malagon et al., 2023).

Replacing extracellular Ca^2+^ with Sr^2+^ greatly enhanced transmission dispersion, producing not only a lot more lingering releases after the stimulation train ends, but also more asynchronous, quantum-sized, supernumerary releases as the GCaMP Sr^2+^ signals took off during stimulation (Figure 8). One possibility is that Sr^2+^ differently interacts with synaptotagmin in the release machinery (Yoshihara and Littleton, 2002; Shin et al., 2003; Tamura et al., 2007), and may also block subtypes of K^+^ channels, causing hyperexcitability of the synaptic terminal membrane (Ganetzky and Wu, 1982, 1983; Ferguson, 1991).

It should be noted that this asynchronous release phenomenon resembles previous reported mutant phenotypes in second messenger-mediated modulatory processes, in particular the cAMP-dependent pathway. Local loose patch-clamp analyses of the mutant phenotypes of the adenylyl cyclase gene *rutabaga* (*rut*) and the phosphodiesterase gene *dunce* (*dnc*) indicate a major role of cAMP-dependent modulation in maintaining release synchronicity (Bukcharaeva et al., 2000; Renger et al, 2000). It is known that cAMP synthesis by adenylyl cyclase is Ca^2+^-dependent through Ca^2+^-Calmodulin binding (Livingstone et al., 1984; Yovell et al., et al., 1992) and hence a strong influence by the local cytosolic Ca^2+^ level.

### Distinctions in plasticity and transmission characteristics of tonic and phasic synaptic boutons – differential excitability and clearance mechanisms

The terms “tonic” and “phasic” initially referred to the contraction kinetics of the slow and fast muscle fibers of crustaceans (Atwood and Raj, 1964). These terms were later extended to describe synaptic terminals displaying distinct modes of short-term plasticity in both crayfish (Kennedy and Takeda, 1965; Msghina et al., 1998, 1999) and *Drosophila* larval (Kurdyak et al., 1994; Lnenicka and Keshishian, 2000; Wu et al., 2015) NMJs. Tonic synapses are known to exhibit facilitation more often (Kurdyak et al., 1994), and thus more suitable for long-lasting slow movement or tonic control, whereas phasic synapses are more prone to depression (Lnenicka and Keshishian, 2000), which are associated with triggering fast, sudden, short-duration movement, resembling the twitch muscles in vertebrates.

As summarized in Figure 12, our data based on individual bouton measurements confirm that compared to tonic synapses, phasic synapses exhibit a greater tendency to depression (Figure 6B), coupled with more potent facilitation (Figure 6A), larger Ca^2+^ dynamics (Figure 5A, B) and more frequent asynchronous transmission (Figures 1, 3, 4, 8). An ultrastructural correlate for the differences in plasticity is that phasic type Is synaptic boutons may contain smaller numbers of both readily releasable and reserved vesicles for their smaller size (Atwood et al., 1993; Jia et al., 1993). However, more elaborate mechanisms also exist. For example, active zones of phasic type Is synapses have tighter ultrastructural appearance and closer coupling with the vesicles (He et al., 2023; see also the diagram in Figure 12A).

Another important distinction is that the tonic type Ib synaptic terminal excitability is apparently more stringently regulated by K^+^ channel repolarization forces to limit Ca^2+^ influx than phasic Is terminals (Figure 12). As mentioned above, perturbations of combined multiple channel blockers or mutations, a single stimulus of can induce a plateau-level, near-maximum GCaMP response in both Is and Ib boutons. This could be achieved in Is boutons after blocking two types of K^+^ channels in several combinations whereas such double insults are insufficient for Ib boutons, which requires a triple insult with combined blockade of certain three K^+^ channel types (Xing and Wu, 2018a).

In addition, previous studies have demonstrated that tonic type Ib synapses possess larger densities of mitochondria (Atwood et al., 1993; Jia et al. 1993), which enables more potent Ca^2+^ sequestration (Xing and Wu, 2018a, b), as well as a greater ATP production for sustaining tonic activities. More abundant ATP could further enhance Ca^2+^ clearance by extrusion through PMCA (Benham et al., 1992; Lnenicka et al., 2006; Xing and Wu, 2018a). Rapid clearance of Ca^2+^ can restrict its accumulation for facilitation and preventing excessive interaction with the vesicular releasing machinery leading to depletion of the releasable pool of vesicles. On the other hand, ATP itself is critical for mobilization of the reserve vesicle pool to replenish releasable vesicles for sustained transmission (Verstreken et al., 2005), which helps explain why type Ib performs better in sustained transmission and more resistance to depression or vesicle depletion (Figure 12C–F).

Besides the above mechanisms, differences in several potential subcellular and molecular activities and interactions may also contribute to the distinctions between type Ib and Is synapses. For example, Ca^2+^-dependent Ca^2+^ channel inactivation for restricting prolonged Ca^2+^ influx, alternative splicing of Ca^2+^ channels for differential expression of functional distinct isoforms (Lee et al., 1999; Chaudhuri et al., 2004; Thalhammer et al., 2017), and Ca^2+^-dependent activation of adenylyl cyclase and phosphodiesterase, can affect the regulation of short-term plasticity (Abrams, 1985; Zhong and Wu, 1991; Renger et al., 2000; Ueda and Wu, 2012, 2015). Such potential enzymatic action and protein modulation steps can be further elucidated by their temporal characteristics and sequence in the functional cascade, in parallel with the status of cytosolic residual Ca^2+^ accumulation and distribution, as indicated by GCaMP signals.

In summary, we report the use of a novel experimental protocol that can help establish the temporal characteristics of GCaMP signals and corresponding short-term plasticity in presynaptic boutons of the *Drosophila* NMJ. This approach may open an avenue to further genetic and pharmacological analyses in this widely adopted neurogenetic model system. It can also help proper interpretation of GCaMP signals in synaptic studies and elucidate the Ca^2+^-dependent molecular mechanisms underlying synaptic facilitation and depression.

## Supporting information

Last page of the manuscript, i.e. List of Supplement Videos

Last page of the manuscript, i.e. List of Supplement Videos

Last page of the manuscript, i.e. List of Supplement Videos

Last page of the manuscript, i.e. List of Supplement Videos

Last page of the manuscript, i.e. List of Supplement Videos

## List of supplement videos

Supplement Video 1. GCaMP imaging of neuromuscular junction (NMJ) in 0.1 mM Ca^2+^. The data corresponds to Figure 1. All video frames have been converted to heat maps. The on- and off-signal of stimulation (40 Hz, 2 s) is indicated by a filled white square in the bottom right corner. The same rule applies to the following videos.

Supplement Video 2. NMJ GCaMP imaging in 0.2 mM Ca^2+^. Corresponds to Figure 2A1 – A4.

Supplement Video 3. NMJ GCaMP imaging in 0.5 mM Ca^2+^. Corresponds to Figure 2C1 – C4.

Supplement Video 4. NMJ GCaMP imaging with 0.5 mM Ca2+ in micropipette, and 0 Ca2+ in bath. The micropipette was on type Ib boutons. Corresponds to Figure 3A1 – A4.

Supplement Video 5. NMJ GCaMP imaging with 0.5 mM Ca2+ in micropipette, and 0 Ca2+ in bath. The micropipette was on type Is boutons. Corresponds to Figure 3 C1 – C4.

